# Machine learning-assisted discovery of growth decision elements by relating bacterial population dynamics to environmental diversity

**DOI:** 10.1101/2022.02.10.479953

**Authors:** Honoka Aida, Takamasa Hashizume, Kazuha Ashino, Bei-Wen Ying

## Abstract

Microorganisms growing in their habitat constitute a complex system. How the individual constituents of the environment contribute to microbial growth remains largely unknown. The present study focused on the contribution of environmental constituents to population dynamics via a high-throughput assay and data-driven analysis of a wild-type *Escherichia coli* strain. A large dataset constituting a total of 12,828 bacterial growth curves with 966 medium combinations, which were composed of 44 pure chemical compounds, was acquired. Machine learning analysis of the big data relating the growth parameters to the medium combinations revealed that the decision-making components for bacterial growth were distinct among various growth phases, e.g., glucose, sulfate and serine for maximum growth, growth rate and growth delay, respectively. Further analyses and simulations indicated that branched-chain amino acids functioned as global coordinators for population dynamics, as well as, a survival strategy of risk diversification to prevent the bacterial population from undergoing extinction.

## Introduction

Highly diversified microorganisms grow in highly differentiated habits^1, 2^. The measurement of diversity in both genetics and the environment is essential to understand community outcomes as an ecological cause and/or consequence^3^ and the evolutionary and responsive strategies constrained by the environment^4, 5^. To date, studies have focused more on genetic diversity, e.g., metagenomics^6, 7^ and microbial communities^8, 9^, than on environmental diversity, despite the high complexity of both microbes and environments^10, 11^. It remains unclear how the individual constituents of the environment (i.e., habitat) contribute to the population dynamics of the microbe or community. Mimicking the environmental diversity in the laboratory by reconstituting the environment (e.g., medium) with known components of defined amounts or magnitudes might be applicable to address this issue.

Microbial population dynamics are commonly represented by growth curves^12, 13, 14^. How a microbial population (species) fits the habitat (environment) has largely been evaluated by three parameters derived from the growth curve, i.e., the lag time, growth rate and saturated population size, which quantitatively represent the lag, exponential and stationary phases of the growth curve, respectively^15, 16^. The lag time has been reported to be crucial for bacterial growth under environmental stress^17, 18^. The growth rate has been associated with proteome allocation^19, 20^, ribosome function^21, 22^ and gene expression^23, 24^; thus, it represents the adaptiveness (fitness) of the microbial population^25, 26^. The three growth parameters are likely coordinated with each other. Previous studies have observed trade-offs between the growth rate and either the population size^27, 28^ or the lag time^29^ within identical species, as well as correlated changes in the growth rate and saturated population size among genetically diversified strains^30, 31^. Whether and how environmental diversity affects the three growth parameters remain unknown.

To address these questions, a quantitatively high-throughput survey linking growth parameters to environmental diversity is required. As the microbial population dynamics have been shown to be strongly dependent on the growth medium^12^, relating the bacterial growth profile to the medium constitution is applicable to address the issue. Recently, both high-throughput technologies for bacterial growth analysis^15^ and data-driven computational approaches have been developed for studying complex systems^32, 33, 34^. In particular, machine learning (ML) techniques have been widely applied to studies on genetic diversity^35, 36^, metabolic engineering^37^ and population dynamics^38, 39, 40^. Combining ML approaches with the use of high-throughput measurements of a well-known microbe in well-defined environments has become practical for comprehensive quantitative evaluation of the contribution of environmental factors (e.g., chemical compositions of the habitats) to microbial population dynamics (e.g., bacterial growth). In the present study, a large dataset describing the bacterial population dynamics in a broad environmental gradient of largely varied combinations was experimentally acquired under well-controlled laboratory conditions. ML prediction and niche broadness analysis of the big data linking bacterial growth to environmental diversity (i.e., medium combinations) were performed. The bacterial growth strategy was investigated by means of data-driven approaches.

## Results

### Relating bacterial growth to environmental diversity

Precise bacterial growth profiling was performed by a high-throughput growth assay in varied medium combinations, which were prepared with 44 pure chemical substances that are commonly used in different microbial culture media (Table S1). As the chemical substances are ionized in solution, these medium combinations finally comprised 41 components (e.g., metal ions, amino acids, etc.) whose concentrations varied broadly on a logarithmic scale (Fig. 1B). In brief, a total of 12,828 growth curves of *E. coli* BW25113 grown in 966 different medium combinations were acquired (Fig. 1A). Three parameters, the lag time (*τ*), maximal growth rate (*r*) and saturated population density (*K*), were subsequently calculated according to the growth curves, which represented the quantitative features in the lag, exponential and stationary growth phases, respectively (Fig. 2A). The averaging of the biological replications and the removal of the unreliable measurements finally resulted in 961, 961 and 937 values of *τ*, *r* and *K*, respectively (Table S2). The three parameters all presented multimodal distributions in response to environmental variation (Fig. 2B), which agreed well with the rugged fitness landscapes proposed for adaptive evolution^41^ and the immune response^42^.

**Figure 1.**
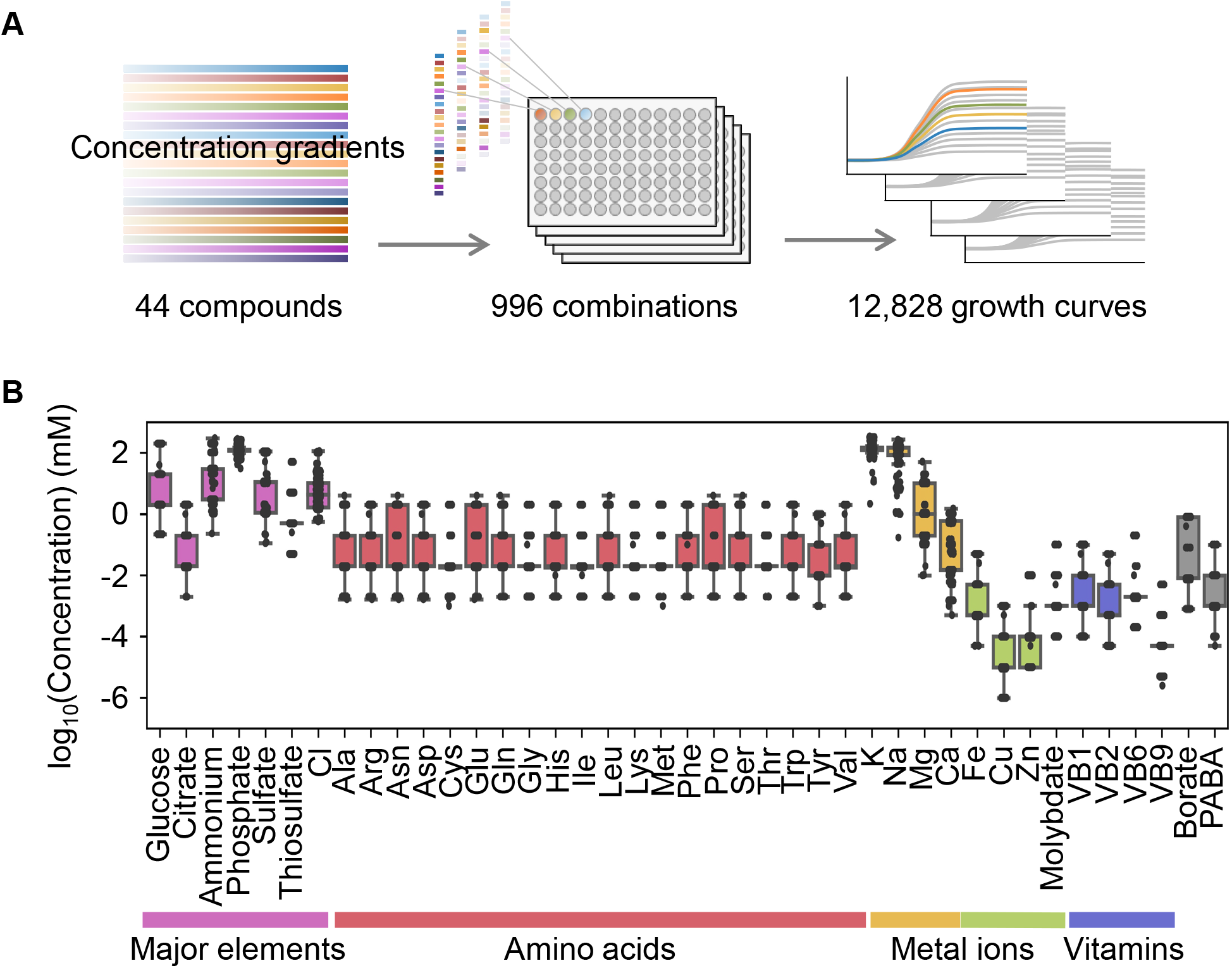
Relating bacterial growth to environmental diversity. **A**. Flowchart of experimental conditions and data attainment. Colour gradation indicates the concentration gradient of the pure chemical compound used in the medium combinations. **B**. Concentration variation of the components comprising the medium combinations. Colour variation indicates the categories of elements. The concentrations are indicated on a logarithmic scale.

**Figure 2.**
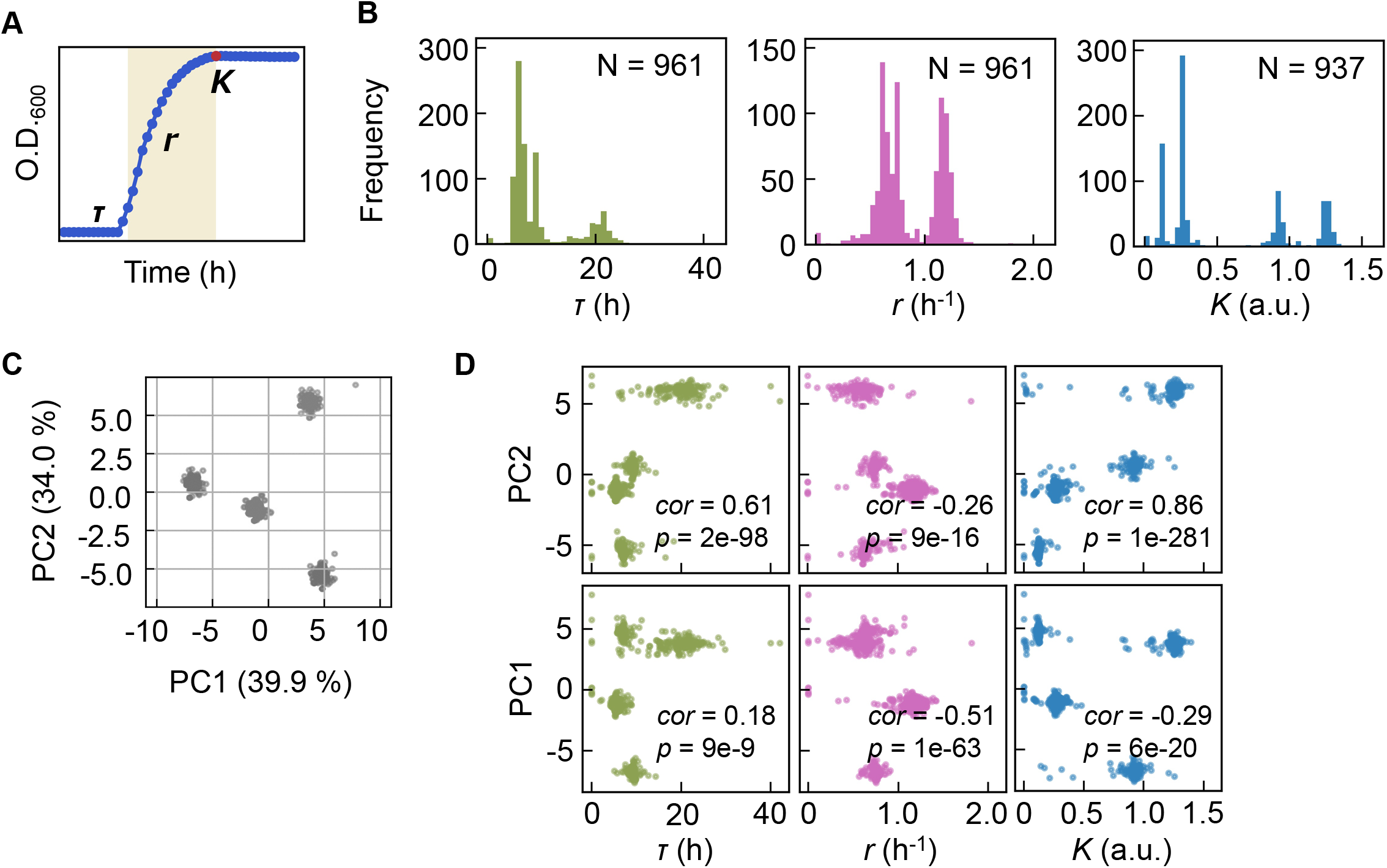
Bacterial growth profiling. **A**. Three growth parameters calculated from growth curves. The lag time (*τ*), the growth rate (*r*) and the saturated population size (*K*) are indicated. **B**. Distributions of the three parameters. The numbers of medium combinations (N) used are indicated. **C**. PCA of medium combinations. The contributions of PC1 and PC2 are shown. **D**. Correlations of the three parameters to the PCs. Spearman’s correlation coefficients and the p values are indicated.

Clustering analyses and principal component analysis (PCA) were applied to the medium combinations. The 966 medium combinations could be mainly divided into four clusters (Fig. 2C), roughly with respect to the multimodality of the distributions (Fig. S1). The three parameters were all correlated with the two main PCs (Fig. 2D), suggesting that bacterial growth was determined by certain common components comprising medium combinations. The results presented an overview of the relationship between the medium combinations and bacterial growth.

### Decision-making components for bacterial growth

Machine learning (ML) approaches were applied to predict the three parameters according to the medium combinations (Fig. 3A). Five representative ML models and an ensemble model were trained and evaluated. The results showed that the prediction accuracy was approximately equivalent among the six ML models (Fig. 3B), independent of the evaluation metrics (Fig. S2). This result implied that the growth parameters could be roughly predicted by varied ML models according to the environmental details, e.g., medium composition. The ML model of the gradient-boosted decision tree (GBDT) was chosen to determine an explainable linkage between bacterial growth and the medium constitution.

**Figure 3.**
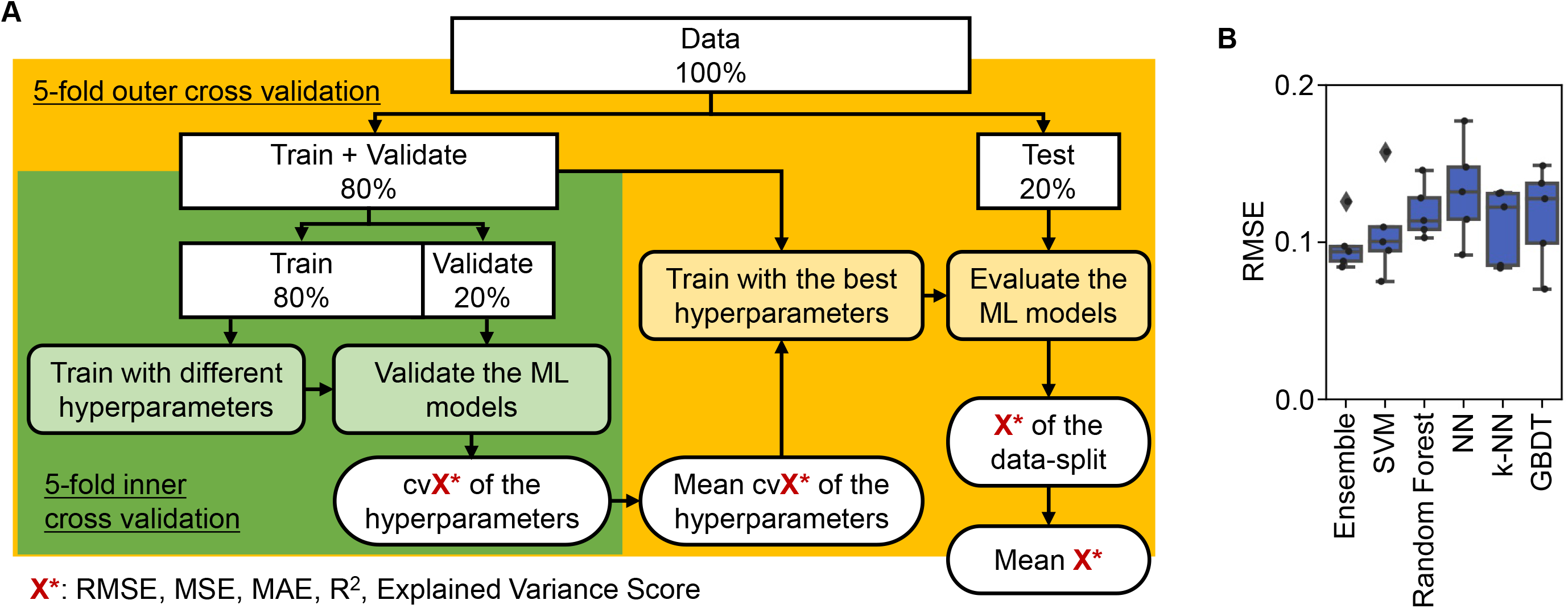
Evaluation of machine learning models. **A**. Workflow of machine learning. **B**. Accuracy of the ML models. Boxplots of the evaluation metrics obtained in the ML prediction of growth rate are shown. The RMSEs of five independent tests are indicated as black points.

Intriguingly, repeated GBDT prediction (Table S3) showed that the growth parameters were largely determined by a single component out of 41 components comprising the medium (Fig. 4A). It seemed that a few key components played a determinant role in bacterial growth, which was supported by the fact that feature reduction in the model training maintained relatively high accuracy (Fig. S3). The top ten features (i.e., components) contributing to the three parameters somehow overlapped (e.g., K, Na and phosphate); nevertheless, the components of the highest priority in governing the three parameters were highly differentiated, that is, serine, sulfate and glucose for *τ*, *r* and *K*, respectively (Fig. 4A). This finding was confirmed by the correlations of the three parameters to the changes in the concentrations of the three components, irrespective of the large variation in other components present (Fig. 4B). It suggested that growth decision-making was highly constrained by a few components and was largely distinguished in response to the growth phase.

**Figure 4.**
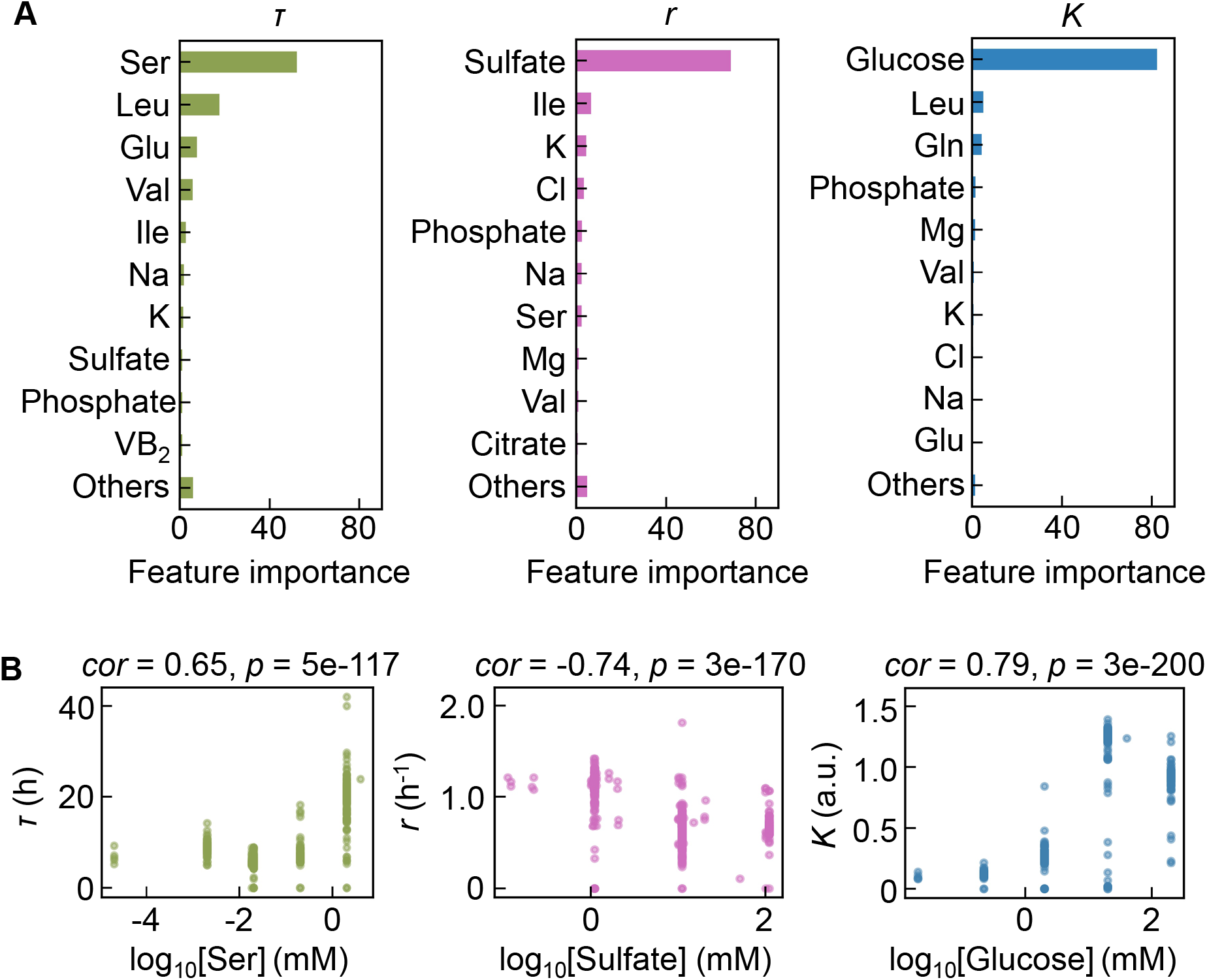
Contribution of the components to bacterial growth. **A**. Relative contributions of the components to the three parameters predicted by GBDT. Ten components with large contributions to the three parameters of *τ*, *r*, and *K* are shown in order. The remaining 31 components are summed as “Others”. **B**. Correlation of the concentrations of the components with the growth parameters. The components with the largest contributions to the three parameters *τ*, *r*, and *K* are shown individually. Spearman’s correlation coefficients and p values are indicated.

### Sensitive components affecting bacterial growth

As the three growth parameters were somehow determinatively decided by a few components, the changes in growth parameters in response to the concentration gradient of each component were evaluated, according to the previous study^43^. Here, the area (i.e., the shadowed space, *S*) above the fitting curve of cubic polynomial regression to the normalized plot was newly defined, in which the maxima of both the concentration gradients and the growth parameters were rescaled to one unit (Fig. 5A). An assortment of fitting curves was acquired for the target component (Figs. S4–S6) because of the various combinations of the remaining 40 components (Fig. 5B). The mean of these *S* values was calculated and designated the sensitivity of the component for bacterial growth in response to the alternative combinations of other components. A larger value of *S* indicated a higher sensitivity of the component, that is, indicated larger changes in the growth parameters due to the variation in the concentration gradients of the other 40 components. Consequently, a total of 41 *S* values were acquired with respect to the three parameters, i.e., *S*_*τ*_, *S*_*r*_ and *S*_*K*_ (Fig. S7, Table S2), which all presented long-tailed distributions (Fig. 5C). The sum of *S*_*τ*_, *S*_*r*_, and *S*_*K*_, which was defined as the global sensitivity (*S_g_*) of the component across the three growth phases, showed a similar long-tailed distribution shape. The four distributions were all likely to follow the power law^44, 45^, which agreed well with the ML-predicted conclusion that only a few components determined the growth. This finding strongly suggested that the fate decision components were present among the 41 components, regardless of the complex interactions among these components.

**Figure 5.**
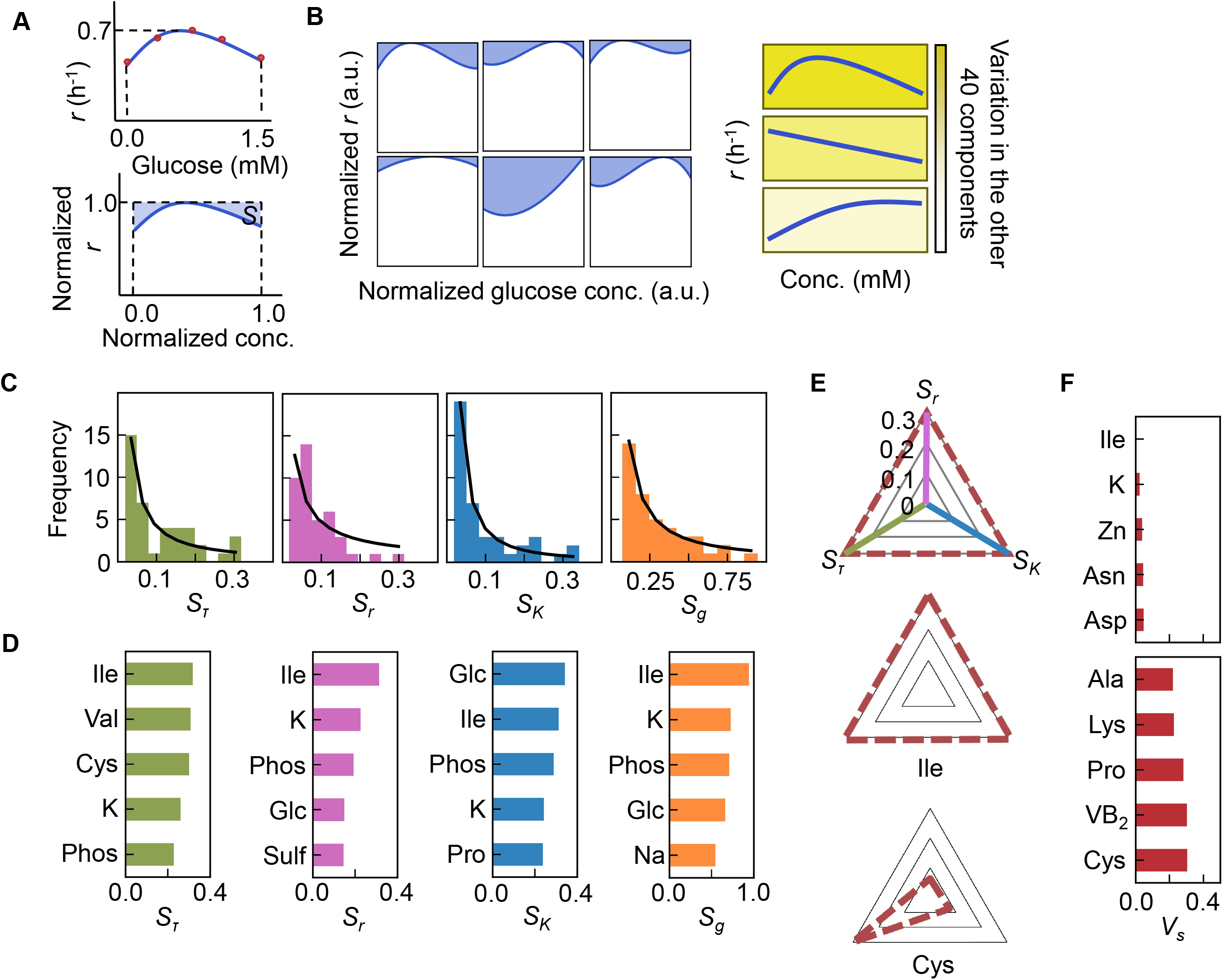
Sensitivity of the components. **A**. Definition of sensitivity. As an example, the upper and bottom panels indicate the regression curve across the concentration gradient of glucose and the normalized regression curve, in which both the concentration gradient and the growth rates are rescaled within one unit, respectively. The shaded area was determined as the sensitivity (*S*) of glucose. **B**. Variation in the sensitivity. Six different regression curves, i.e., six different *S* values, of glucose are shown, which result from the alternative combinations of the other 40 components (left panels). The yellow gradation and blue lines represent the variation in medium combinations and the corresponding regression curves, respectively (right panels). **C**. Distributions of the mean sensitivities. The mean *S* values evaluated according to *τ*, *r*, and *K* are shown as *S*_*τ*_, *S*_*r*_ and *S*_*K*_, respectively. The sum of the three *S* values is shown as *S*_*g*_. The black lines indicate the fitting curves of the power law. **D**. Most sensitive components. The components with the largest *S* values are shown in the order of value. **E**. Balance of sensitivity. The balance of sensitivity is visualized by the triangle of *S*_*r*_, *S*_*K*_ and *S*_*τ*_ in red dotted lines. The solid lines in pink, blue and green represent *S*_*r*_, *S*_*K*_ and *S*_*τ*_, respectively. Those close to or far from an equilateral triangle are determined as the balanced (Ile) or biased (Cys) sensitivity in response to the growth phases, respectively. **F**. Variance of sensitivity. The components with either the smallest or the largest *V*_*s*_ are shown in the order of value. Five components of either balanced or biased sensitivity are shown.

The components with the largest *S* values, i.e., Ile, K, and phosphate (Fig. 5D), overlapped among the three parameters, suggesting that these components were highly sensitive to the fluctuation of other components for all growth phases. In particular, Ile was the most sensitive component, which supported the finding that the components dominating the three second-priority parameters were Leu and Ile (Fig. 4A). Additionally, analysing the variance (*V*_*s*_) of the *S*_*τ*_, *S*_*r*_ and *S*_*K*_ values showed that the smallest and largest *V*_*s*_ values were in Ile and Cys, (Fig. 5E, Fig. S7), indicating general and biased sensitivity for the three growth phases. This finding was independent of the methods used for the evaluation of the variance (Table S4). Overall, Ile and/or branched-chain amino acids (BCAAs) participate commonly in all growth phases and are probably global coordinators for bacterial growth.

### Risk diversification strategy for population survival

The three components serine, sulfate and glucose, determining the growth lag, growth rate and growth yield, respectively (Fig. 6A), could be categorized into the three major elements of nitrogen (N), sulfur (S) and carbon (C). The contribution and mechanism of C and N to population dynamics have been intensively studied^46, 47, 48, 49^, whereas little is known concerning S. To link sulfate to growth, flux balance analysis (FBA)^50^ simulation was performed. The result showed that a decreased growth rate was associated with an increased rate of sulfate uptake (Fig. 6B), supporting the determinative contribution of S to the growth rate. Nevertheless, the FBA simulation did not provide a perfect explanation, as the concentration of sulfate used in the simulation was somehow excessive compared to that used for culture in general. The determinative role of S in the growth rate could result from its function as a material because S is not only the major element in organisms but also a major constituent of the earth (Table S5).

**Figure 6.**
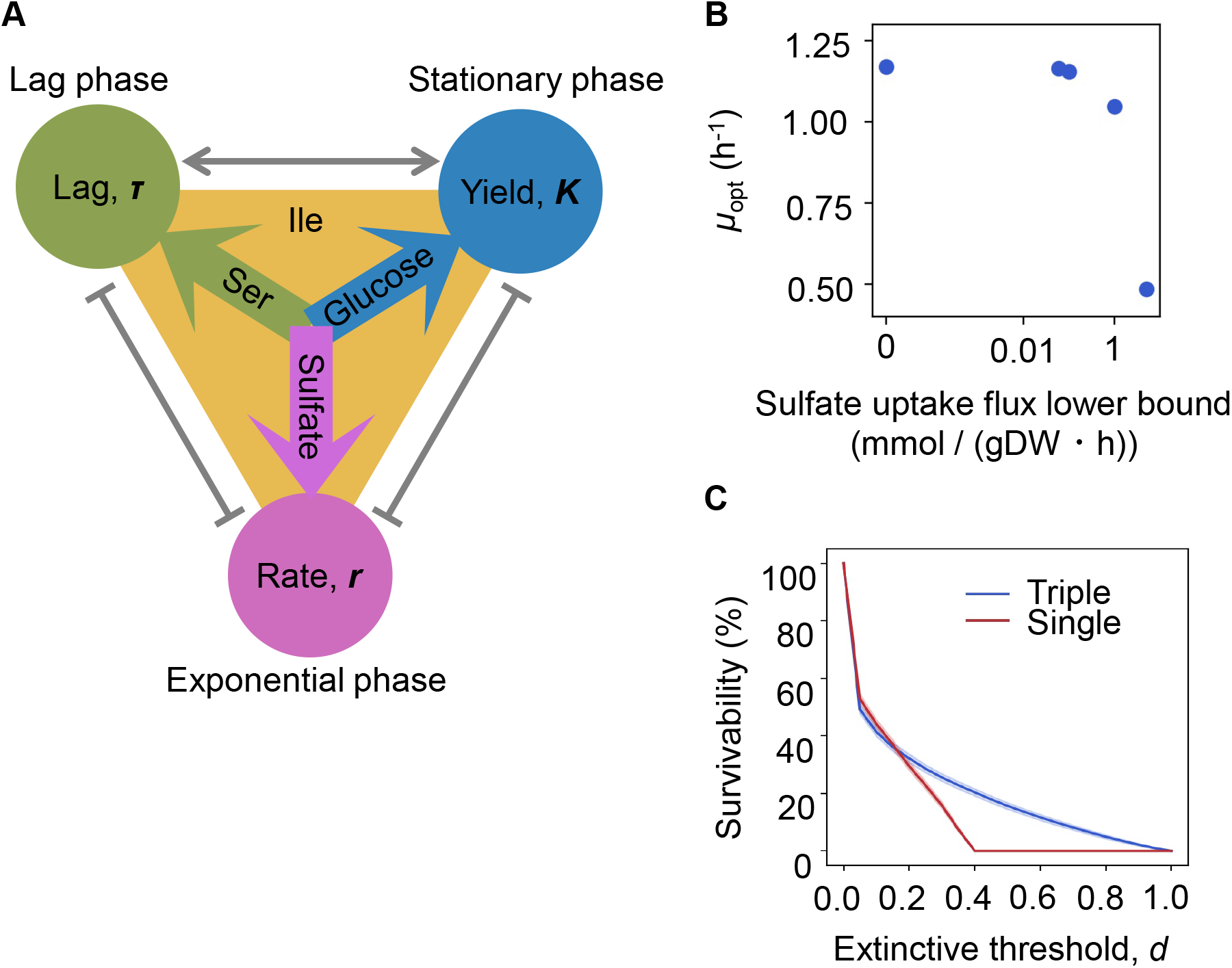
Growth strategy of risk diversification. **A**. Schematic drawing of the decision-making components for bacterial growth. **B**. FBA simulation. The predicted growth rates are plotted against the input rate of sulfate uptake. **C**. Theoretical simulation of survival probability. The blue and red lines represent the growth strategies of the multiple and single decision-makers, respectively. The shading covering the red and blue lines indicates the standard deviation.

Notably, the different elements regulating various growth phases strongly implied risk diversification in fate decisions as a survival strategy. To demonstrate whether the differentiation of elements for growth decisions are a practicable survival strategy, theoretical simulations based on either a single or multiple determinants for the three parameters were additionally performed. Every 1,000 simulations were conducted at the varied threshold (*d*), i.e., the ratio was defined as population distinction. The results showed that a three-component set of decision-makers led to a higher probability of survival, particularly when reducing the extinction threshold (Fig. 6C). The differentiation in fate decision-makers prevented the bacterial population from undergoing extinction, which must be beneficial for the bacteria growing in a fluctuating environment, as it agreed well with the prospected Y-A-S strategy of the microorganisms in nature, i.e., the growth strategy for high yield, resource acquisition and stress tolerance, respectively^51^.

### Coordination in bacterial population dynamics

As the differentiation in decision-making components allowed the independent decision for varied growth phases, the previously reported correlated changes in the growth parameters^27, 28, 29, 30, 31^ were supposed to be weakened. However, the three parameters remained significantly correlated (Fig. 7), which indicated that the risk diversification strategy did not disturb the trade-off or coordination, e.g., *K/r* selection^52^. The correlated changes of the growth parameters might be due to the global participation of Ile and/or BCAAs, as the decision-makers and common sensors. The frequency of Ile and BCAAs coded into the proteins in growing cells was evaluated (Fig. 8A). The relative abundance of amino acids (AAs) was determined as the ratio of the frequency of the target amino acid to the sum of all 20 amino acids in all proteins. Taking into account the variation in the copy number of proteins in growing cells, the frequency of each AA was normalized based on the relative abundance of gene expression (Fig. 8B). The relative expression level of each gene (protein) was calculated as the mean of biologically repeated transcripts, according to previous reports^30, 53^. The results showed that the relative abundances of intracellular Ile and BCAAs were significantly higher than their theoretical ratios, i.e., one or three out of 20 amino acids, 5 or 15%, respectively (Fig. 8C). Although the most abundant AA was not Ile but Leu (Fig. S8), their regulation and metabolism are closely related^54^. The results revealed that the protein building blocks required more BCAAs than other AAs, except Ala and Gly (Fig. S8). The coordination among the three growth parameters might be balanced by BCAAs.

**Figure 7.**
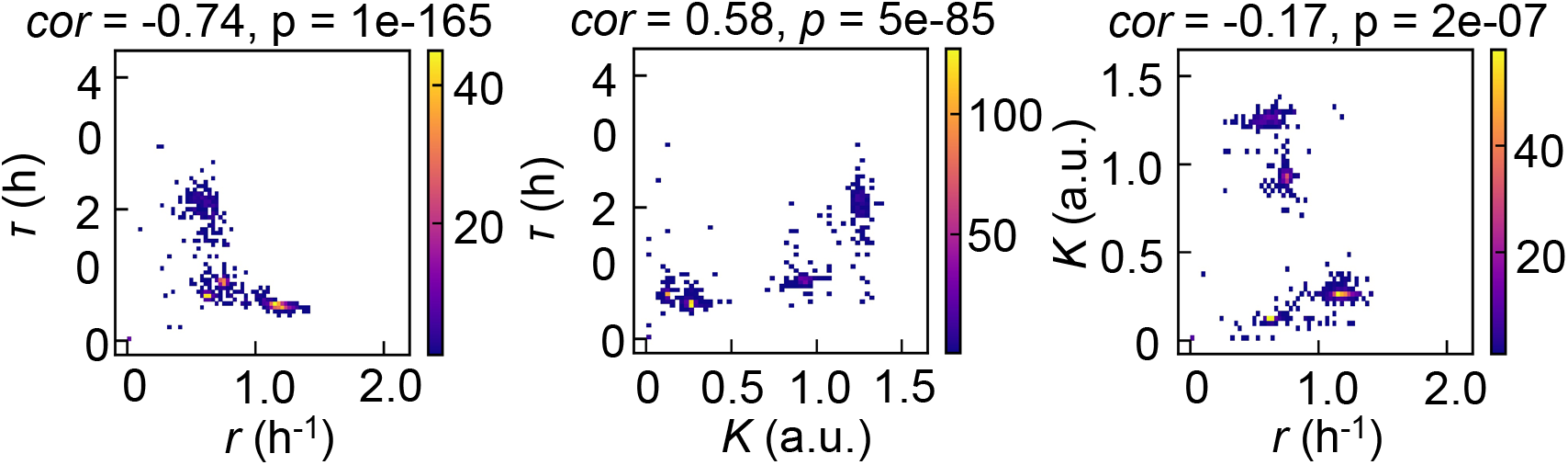
Density plots of the three parameters. Pairs of the three parameters *τ*, *r*, and *K* are plotted as dots. The colour bars indicate the numbers of data points. Spearman’s correlation coefficients and the p values are indicated.

**Figure 8.**
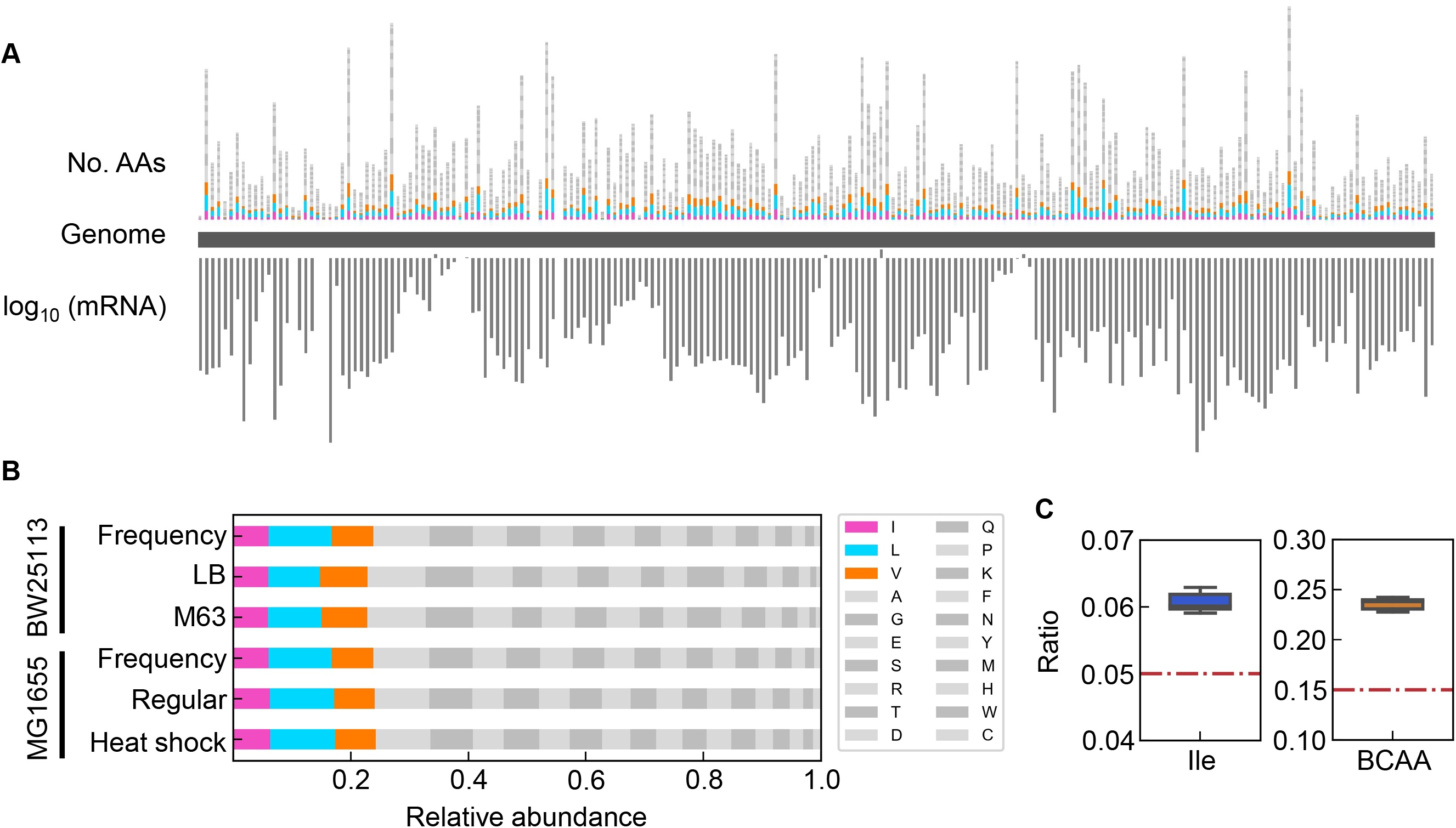
Relative abundance of BCAAs in growing *E. coli*. **A**. Chromosomal distribution of 20 amino acids. The numbers of 20 AAs coded into the proteins are indicated with the upward vertical bars, at the chromosomal positions of the corresponding genes. BCAAs and the remaining 17 AAs are in colour and monotone, respectively. The gene expression level in a logarithmic scale is indicated by the downward vertical bar in grey. **B**. Relative abundance of 20 amino acids. Twenty amino acids are shown with a single letter abbreviation. BCAAs are highlighted. The *E. coli* strains BW25113 and MG1655 are indicated. Frequency represents the relative abundance of the amino acids, while all proteins encoded on the genome are of equivalent amount. LB, M63, regular and heat shock indicate the relative abundance of the amino acids according to the transcriptomes of the *E. coli* cells grown in LB, in M63, at 37 °C and at heat shock conditions, respectively. **C**. Relative abundance of Ile and BCAAs in growing *E. coli*. The boxplots represent the relative ratios estimated according to the genome and transcriptome information, and the red lines indicate the theoretical ratios of the 20 amino acids.

## Discussion

The experimental restriction of the present survey was supposed to be the variety of environmental conditions and the interval of the concentration gradient. First, the study focused on the contribution of chemical conditions to bacterial growth, which was also affected by other environmental conditions, such as temperature and oxygen^55, 56, 57^. As the quantitative adjustment of these conditions was currently impracticable for the high-throughput assay, these conditions were beyond the scope of the present study. The findings on the contribution of medium components to bacterial growth were true under laboratory conditions. Second, the concentration gradients of the components were prepared as broadly as possible to achieve the maximal solubility available for medium combinations, some of which were largely different from those used in the laboratory and/or found in nature^58, 59^. The broad range of concentrations allowed us to acquire a boundless fitness landscape across the greatest environmental gradient and led to a wide concentration interval, i.e., changes on a logarithmic scale, in the growth assay. The concentration gradient of the most sensitive range to change bacterial growth might have been missed or masked. As the proper range of concentration gradients for sensitive growth change remained a black box, practicable conditions were applied. Theoretically, the issue concerning the best concentration gradient could be solved by extensive growth assays associated with the ML prediction of concentration determination (rescale), that is, introducing the predicted concentration to the following round of the growth assay and repeating the ML training and the experimental test. The extended repeats would result in the best medium combination for bacterial growth, which could be applied for culture optimization and development.

Since the accuracy and reliability of ML are largely dependent on the quality and quantity of the training data, the impact of the experimental data on the ML models was carefully assessed. First, the representative ML models were compared (Fig. 3, Fig. S2). The best accuracy was acquired with the ensemble model (Fig. S2); nevertheless, as it required the longest time for model training (Fig. S9) and was uninterpretable, the GBDT model was finally employed. Second, whether the abundance of the dataset affected the accuracy of the GBDT model was evaluated. The amount of data used for model training varied from 10%~90% of the entire big data set. Although a small amount of data (~10%) led to a relatively high accuracy on average, the variance in the accuracy of repeated model training was too large to reach a reliable prediction (Fig. S10). An increase in the abundance of the training data decreased the variance of the model accuracy (Fig. S10), demonstrating that a sufficiently large dataset was essentially required to achieve robust ML prediction for the biological experiments, as discussed in the different ML-associated microbial studies^60^. The dataset used here was large enough to grant a small variance, indicating the robust result of model training. Third, whether the accuracy of prediction was attributed to the experimental errors was evaluated. An equal amount of training data (n = 400) with varied experimental accuracy, i.e., the variance of biological replication (*CV* = 0.05~0.12), was used to test the accuracy of the ML models. Intriguingly, the training data with large variance, i.e., large experimental error caused by biological replication, resulted in the high accuracy of ML, in comparison to those with small variance, which led to the decreased accuracy of ML (Fig. S11). Accordingly, the entire experimental dataset, regardless of the experimental error, was used for ML to draw the conclusion presented here.

The multimodal distributions of the three parameters (Fig. 2A) reflected a broad variation in medium combinations; nevertheless, whether the main conclusion regarding the differentiated decision-makers for varied growth phases was biased by the medium combinations was evaluated. First, the variation in the concentration of each component was counted. The most abundant variation of concentration was that for the chlorine ion (Cl), which was a low-priority contributor to growth, whereas the decision-making components showed either high or low variation of concentrations, such as for sulfate or glucose, respectively (Table S6). Second, although the amino acids presented equivalent variations in concentration, only Ile, Ser and Leu were determined to be the growth determinative components. Finally, even if the multimodal distributions of the three parameters were arbitrarily divided into two monomodal-like distributions for data separation, which led to the reduced abundancy of the dataset, the differentiation in decision-making components for the three growth phases remained (Figs. S12–S14). As the data separation reduced the variety of medium combinations, the highest-priority components were either similar or varied from those identified while using the whole dataset (Fig. 4). This result indicated that the diversity of experimental conditions, i.e., the abundance of training data, could influence the ML prediction. The present study applied the exceeding range of the concentration gradient and the high variability of medium combinations, which might cover the landscape of population dynamics as broadly as possible in the laboratory; therefore, the finding of the differentiation in the components deciding the three growth phases was independent of the experimental restriction.

In summary, the present study provided an informative and quantitative big data set relating bacterial growth (population dynamics) to environmental factors and valuable knowledge for introducing data science to biological experiments. The differentiation in growth decision-making components in the lag, exponential and stationary phases protected the bacterial population against extinction. This finding revealed a common and simple strategy of risk diversification for bacterial growth in conditions of excessive resources or starvation, which is a reasonable approach in evolution and ecology. As a successful example, the present study demonstrated that investigating the microbial world by data-driven approaches allows us to perceive highly intriguing insights that were inconceivable by traditional experimental approaches.

## Materials and Methods

### Bacterial strain and stock preparation

The wild-type *E. coli* strain BW25113 was used, which was provided by the National BioResource Project (National Institute of Genetics, Shizuoka, Japan). To reduce the experimental errors of the repeated growth assay on different days, common stocks of the exponentially growing *E. coli* cell culture were prepared beforehand, as described previously^61^. In brief, the *E. coli* cells were cultured in 5 ml of M63 minimal medium using a bioshaker (BR23-PF, Taitec) at 200 rpm and 37 °C. The cell culture was stopped when its optical density measured at 600 nm (OD_600_) reached ~ 0.1. The culture was subsequently divided into a small portion (60 μL) in 1.5 mL microtubes (Watson) and stored at −80 °C for future use. Hundreds of aliquots (stocks) were prepared at once and disposably used in the growth assay; that is, aliquots were used only once, and remaining cultures were discarded.

### Medium composition and combinations

A total of 44 pure chemical substances, determined according to the literature^62, 63^, were all commercially available (Wako or Sigma). The minimal concentrations of these compounds were set at zero in general, and the maximal concentrations were determined individually according to the literature or laboratory manuals (Table S1). In addition, the concentrations of the compounds rarely used in the known media were experimentally examined (Figs. S15 and S16). According to the determined maximal concentration, stock solutions of these chemical substances were prepared in advance for the easy preparation of medium combinations. The chemical substrates were dissolved in highly pure water (Direct-Q UV, Merck) at high concentrations. Subsequently, the resultant solutions were sterilized, either using a sterile syringe filter with a 0.22 μm pore size and hydrophilic PVDF membrane (Merck) or by autoclaving at 121 °C for 20 min. The stock solutions were divided into aliquots (10~100 μL) in 1.5-mL microtubes (Watson) and stored at −30 °C for future use. A total of 100~300 stocks were prepared at once for individual chemical substrates. To avoid repeated thawing and freezing of the stock solutions, aliquots were used only once. The medium combinations were prepared by mixing the stock solutions (aliquots) just before the growth assay. The concentrations of the substrates were varied on a logarithmic scale, and only a single substrate was altered for each assay. A total of 966 combinations were tested in the growth assay (Table S2).

### Growth assay

The high-throughput growth assay was conducted to acquire the growth curves in the medium combinations, as described previously^40^. The culture stocks were diluted 1,000-fold with 5 ml of fresh media of varying medium combinations in 5 ml tubes (Watson). The diluted cell culture mixtures were loaded into a 96-well microplate (Costar) in four-to-six wells (200 μl per well) with varied locations per medium combination. The 96-well plates were incubated in a plate reader (Epoch2, BioTek) with a rotation rate of 567 rpm at 37 °C. The temporal growth of the *E. coli* cells was detected at an absorbance of 600 nm, and readings were obtained at 30-min intervals for 24 to 48 h. A total of 12,828 reliable growth curves were acquired.

### Data processing and calculation of the growth parameters

The temporal OD600 reads were exported from the plate reader and processed with Python, as described in detail elsewhere^40^. The growth parameters *τ*, *r* and *K* were evaluated according to previous reports^40, 64^ using a previously developed Python program^40^. In brief, *τ* was determined as the time when the increase in OD600 was observed in five consecutive reads; *r* was defined as the mean of three continuous logarithmic slopes of every two neighbouring OD600 values within the exponential growth phase using “gradient” in the “numpy” library; and *K* was calculated as the mean of three continuous OD600 values including the maximum, which was determined using “argmax” in the “numpy” library.

### Principal component analysis and clustering

Principal component analysis (PCA)^65, 66^ was performed using “PCA” in the “decomposition” module from “scikit-learn”^67^. The concentrations of 41 components were normalized within one unit, and the 966 combinations were used as input. The principal component scores of PC1 and PC2 were used for the correlation analysis of the three growth parameters. Clustering of the PC1-PC2 scores was performed using “KMeans” in the “cluster” module of the “scikit-learn” library.

### Machine learning models and evaluation

Machine learning was performed using a supercomputer, the Cygnus system (NEC LX 124Rh-4G). The machine learning models of gradient-boosted decision tree (GBDT), k-nearest neighbour (k-NN), neural network (NN), random forest and support vector machine (SVM) were performed using “GradientBoostingRegressor” in the “ensemble” module, “KNeighborsRegressor” in the “neighbors” module, “MLPRegressor” in the “neural_network” module, “RandomForestRegressor” in the “ensemble” module and “SVR” in the “svm” module, respectively. The ensemble model was performed using “StackingRegressor” in the “ensemble” module and “LinearRegression” in the “linear_model” module. Data normalization was performed using “StandardScaler” in the “preprocessing” module for k-NN and NN and “MinMaxScaler” in the “svm” module for SVM. All these modules were in the “scikit-learn” library.

A fivefold nested cross validation was performed to evaluate the ML models. A grid search was used for the hyperparameter search, as follows. In the GBDT model, “random_state” and “n_estimators” were configured as 0 and 300, respectively; “learning_rate” and “max_depth” were searched from 0.001 to 0.5 in increments of 0.005 and among 2, 3, 4, and 5, respectively. In the k-NN model, “n_neighbors” was searched among 1, 2, 3 and 4. In the NN model, “solver” and “alpha” were configured as “adam” and 0.001, respectively; “hidden_layer_sizes” was searched among (100,100,100), (100,100), (50,50) and (50,50,50). In the random forest model, “random_state” and “n_estimators” were configured as 0 and 300, respectively; “max_depth” was searched among 2,3 and 4. In the SVM model, the “kernel” was configured as “rbf”; “C”, “gamma” and “epsilon” were searched from 2^−5^ to 2^10^, 2^−20^ to 2^10^ and 2^−10^ to 2°, respectively, in increments of 2^2^. All other hyperparameters were used as default.

The metrics adopted to estimate the accuracy of the ML models were determined as follows. The coefficient of determination (R^2^), mean squared error (MSE), mean absolute error (MAE), and explained variance score were calculated using “r2_score”, “mean_squared_error”, “mean_absolute_error”, and “explained_variance_score” in the “metrics” module of the “sklearn” library, respectively. The root mean squared error (RMSE) was calculated with the MSE values using “sqrt” in the “numpy” library.

### GBDT prediction and feature reduction

A regression model was created by using the log-transformed concentrations of the components. The “feature_importances_” attribute represents the importance of each component to the creation of the model. Outer and inner cross validation was performed using “cross_val_score” in the “model_selection” module of the “scikit-learn” library. The hyperparameters were searched using “GridSearchCV” in the “model_selection” module of the “scikit-learn” library. “learning_rate” and “max_depth” were searched from 0.01 to 0.5 in increments of 0.01 and among 2, 3, 4 and 5, respectively. “n_estimatiors” was configured at 300, and the other hyperparameters were set to default values. The “feature_inportance_” values were calculated by fivefold cross validation, and the mean of the five values was used as the result of the GBDT prediction. Feature reduction was performed using “RFE” in the “feature_selection” module of the “scikit-learn” library, in which the “estimator” of the “RFE” was the GBDT model.

### Evaluation of sensitivity

The changes in the growth parameters associated with the concentration gradient of each component were evaluated by curve fitting of a cubic polynomial as described previously ^43^.

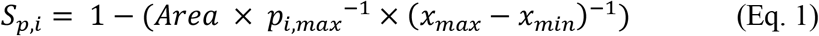

Here, *S*_*p,i*_, *Area*, *p*_*i, max*_, *x*_*min*_ and *x*_*max*_ represent the sensitivity evaluated with any of the growth parameters in Condition *i*, the area under the regression curve, the largest value of the growth parameter in Condition *i*, and the minimum and maximum concentrations of each chemical component, respectively. The sensitivity was further evaluated as follows.

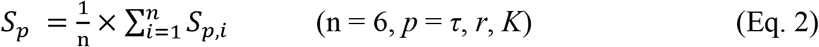

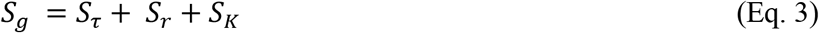

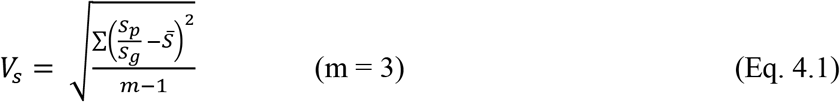

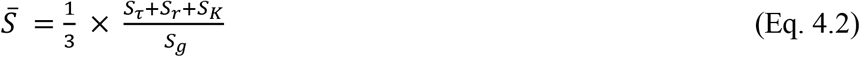

Here, *S*_*τ*_, *S*_*r*_ and *S*_*K*_ represent the sensitivity of *τ*, *r* and *K*, respectively. *S*_*g*_ and *V*_*s*_ represent the sum and the variance of *S*_*τ*_, *S*_*r*_ and *S*_*K*_, respectively. Additionally, four different methods were applied to estimate *V*_*s*_, as follows.

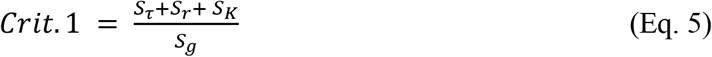

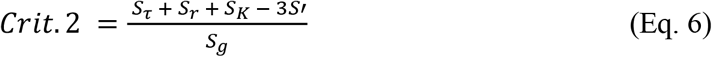

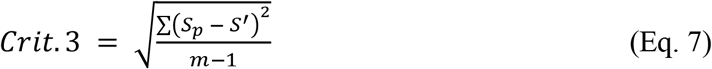

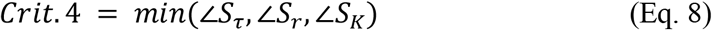

Here, *S’* indicates the mean of *S*_*τ*_, *S*_*r*_ and *S*_*K*_. ∠*S*_*τ*_, ∠*S*_*r*_, and ∠*S*_*K*_ represent the three angles calculated from the triangle (Fig. S8).

### Flux balance analysis simulation

Flux balance analysis (FBA) simulation was performed using the open software COBRAme^68^. iJL1678b-ME and qMINOS, which were available in the Docker images^69^, were used as the model and the solver, respectively, where “mumax” and “precision” were set as 2 and 1E-6, respectively. Four out of 41 components, i.e., VB9, VB2, borate and PABA, were excluded in the simulation, as they were absent in the ME model. The lower bounds of the efflux of the components were set as the negative values, which allowed the *E. coli* cells to take them up from the media. The lower bound of the efflux of amino acids and citrate were set to −10, and −1000 was set for the others. The lower bounds of the efflux of selenite, selenite, tungstate, Li, Sc, and Tl were set to zero, and those of cobalt, Mn, Ni, RNase_m5, RNase_16, and RNase_m23 were set as −0.00001, −0.001, −0.001, −1, −1 and −1, respectively, because these components were absent in the present study. The uptake of sulfate was fixed by setting the upper bound of the efflux to a negative value to predict the growth rate when the uptake of sulfate was varied.

### Genomic datasets and annotation

The genome and transcriptome datasets of the *E. coli* BW25113 and MG1655 strains were obtained from GenBank (CP009273 and NC_000913) and GEO (GSE33212 and GSE136101), respectively. The gene (protein) annotation and counting of the amino acids were processed using BioPython^70^.

### Theoretical simulation of survival probability

The simulation of population dynamics over 24 hours was conducted according to the following equations.

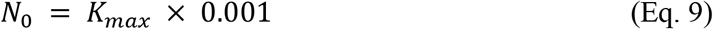

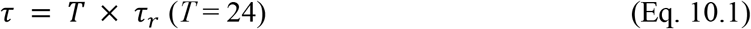

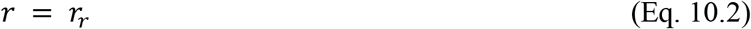

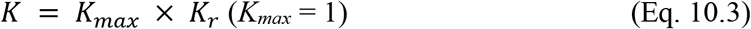

Here, *N_0_*, *K_max_ τ_r_*, *r*_*r*_, and *K*_*r*_ are the initial population, the population maximum, and the three variables *τ*, *r* and *K*, respectively. In the case of triple independent decision-makers, the values of *τ_r_*, *r*_*r*_, and *K*_*r*_ were randomly selected from 0 to 1 without coordinated change. In the case of a single common decision-maker, once any of the three parameters was randomly selected from 0 to 1, the other two were decided as follows.

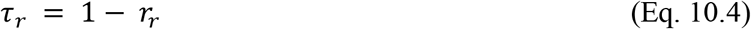

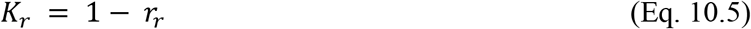

The population dynamics were defined as follows.

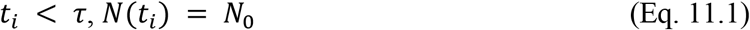

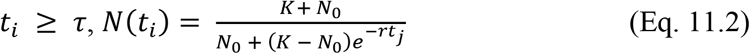

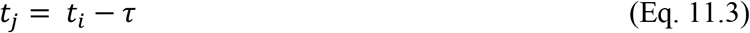

where *N*(*t*_*i*_), *t*_*j*_ and *t*_*i*_ are the population size at time *t*_*i*_, any time point within the exponential phase and any time point from 0 to 24 h in a 0.5 h interval, respectively. Whether the population was extinct or survived was determined according to the survival threshold, *d*, as follows.

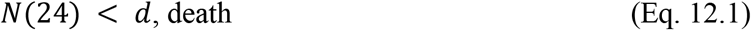

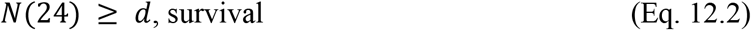

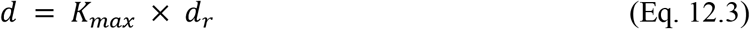

Here, *N*(24) and *d*_*r*_ are the final population size at 24 h and the threshold varying from 0 to 1 in increments of 0.05, respectively. The survival probability was defined as the frequency of “survival” in every 1,000 simulations at each *d*_*r*_.

### Separation of the multimodal distributions

Gaussian kernel density estimation was used to determine the boundaries of the multimodal distributions, which were considered bimodal, for data separation of the growth parameters. The probability density function was conducted using “gaussian_kde” in the “stats” module of the “scipy” library, in which “bw_method” was configured as 0.3. These distributions were divided vertically into 1,000 equal areas. The trough point, i.e., the smallest area, for data separation was determined using “argrelmin” in the “signal” module of the “scipy” library. The three growth parameters were independently divided into two datasets of low and high mean values. The following GBDT prediction of *τ*, *r* and *K* was performed on the three separated datasets (Figs. S12–S14).

## Acknowledgements

We thank NBRP (Japan) for providing the *E. coli* strain. This work was supported by a JSPS KAKENHI Grant-in-Aid for Challenging Exploratory Research (grant number 21K19815) and partially by a JSPS KAKENHI Grant-in-Aid for Scientific Research (B) (grant number 19H03215).

## Author contributions

BWY conceived the research; HA and KA performed the experiments; HA, MH and BWY analysed the data; HA drafted the manuscript and graphics; BWY rewrote the paper; and all the authors approved the paper.

## Supplementary Figures

**Figure S1.**
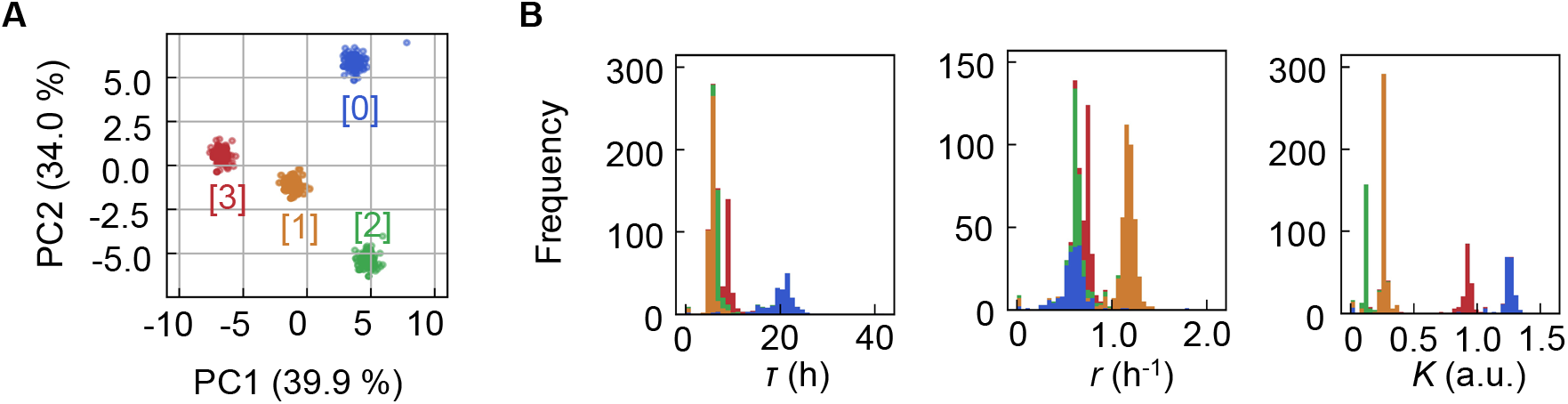
Clustering of the medium combinations. **A**. Clusters of medium combinations. **B**. Distributions of *τ*, *r*, and *K* coloured by the four clusters.

**Figure S2.**
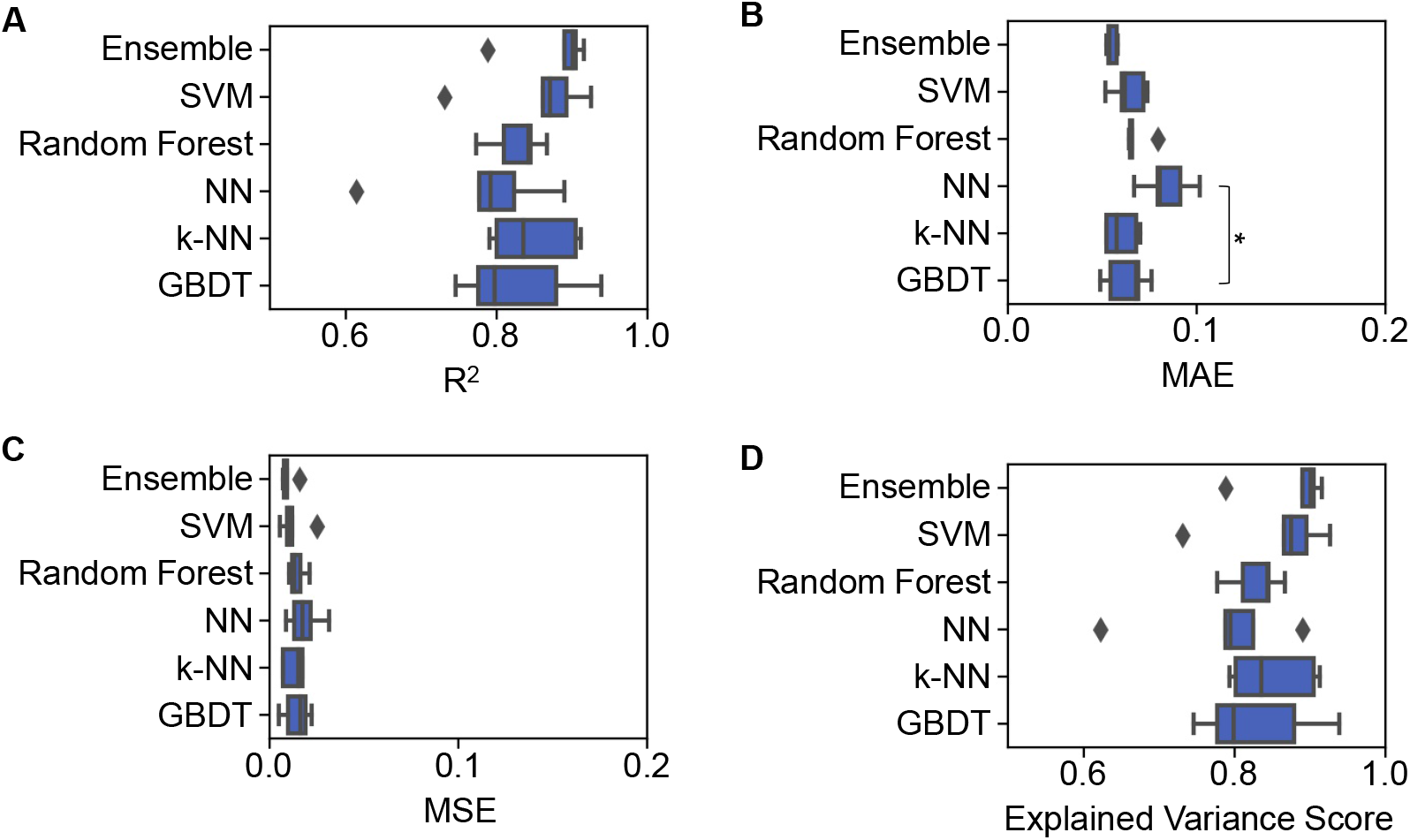
Accuracy of the ML models. Boxplots of four different evaluation metrics, i.e., R^2^ (**A**), MAE (**B**), MSE (**C**) and explained variance score (**D**), obtained in the ML prediction of the growth rate are shown. The results from 5 independent tests are indicated as black points. Asterisks indicate statistical significance (p < 0.05).

**Figure S3.**
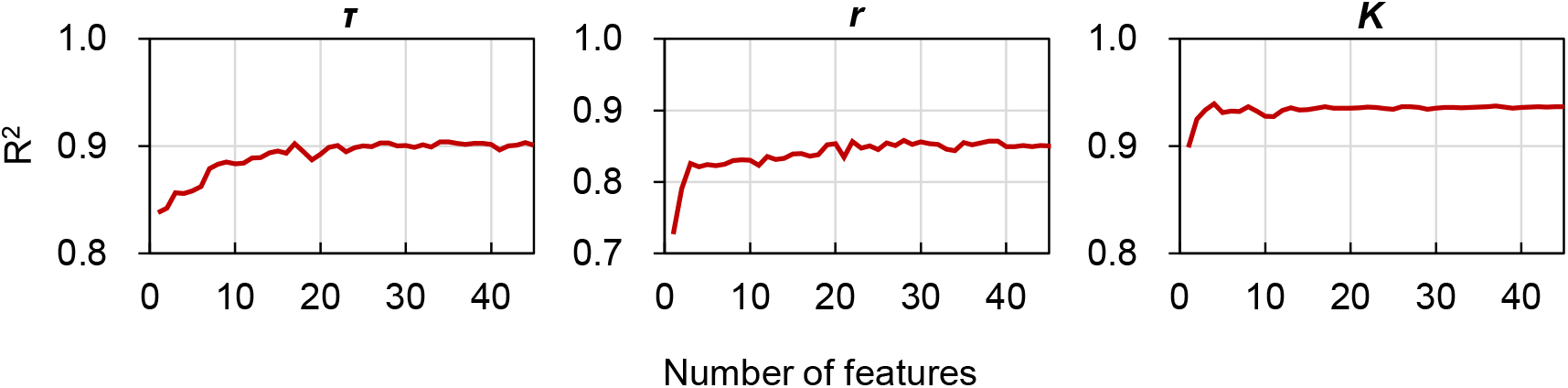
Feature reduction in GBDT. The accuracy of the GBDT prediction using one to 41 features (components) is shown. The three parameters *τ*, *r*, and *K* are indicated.

**Figure S4.**
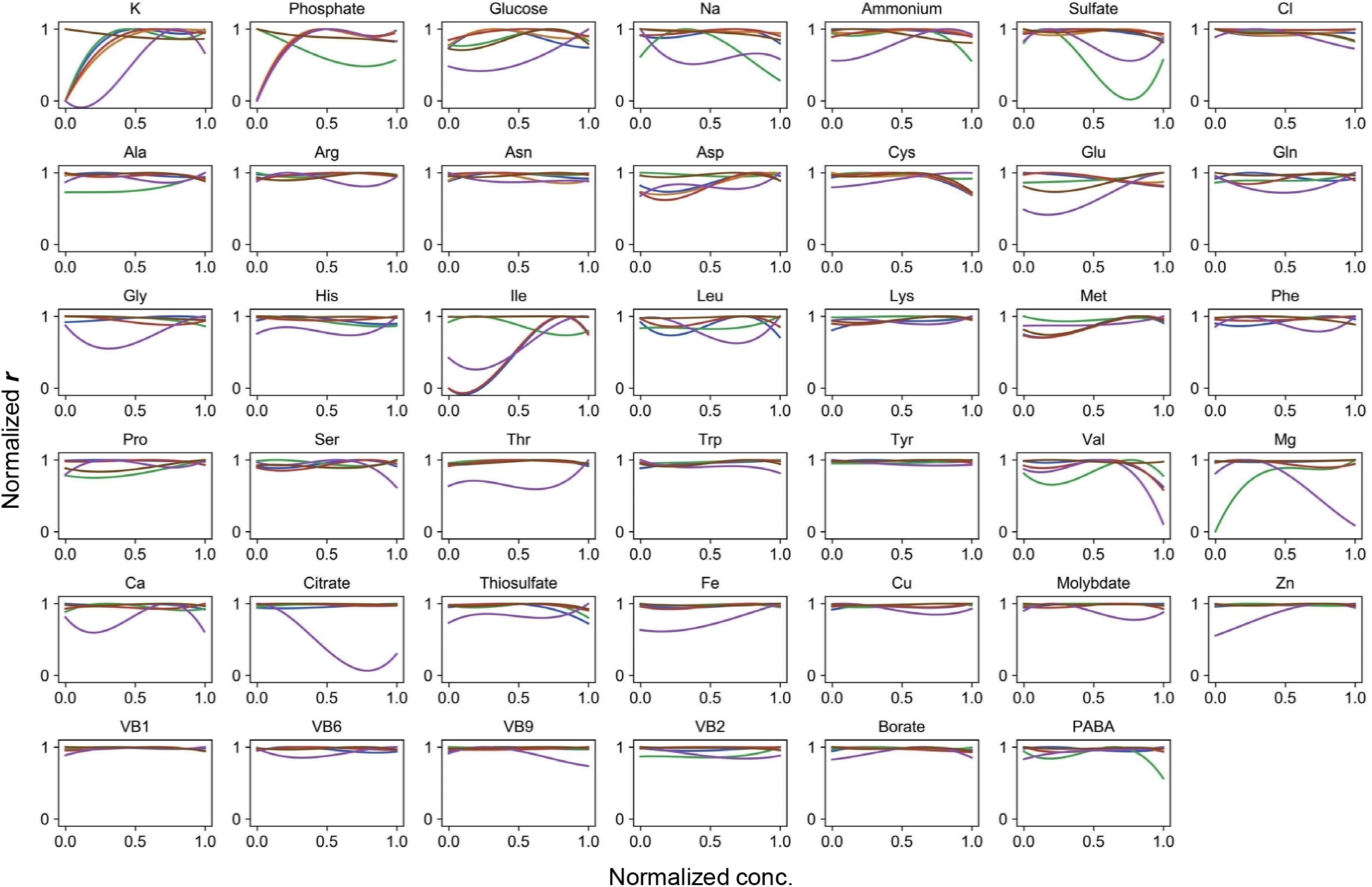
Normalized regression curves of the growth rates. Colour variation indicates the alternative combinations of the other 40 components.

**Figure S5.**
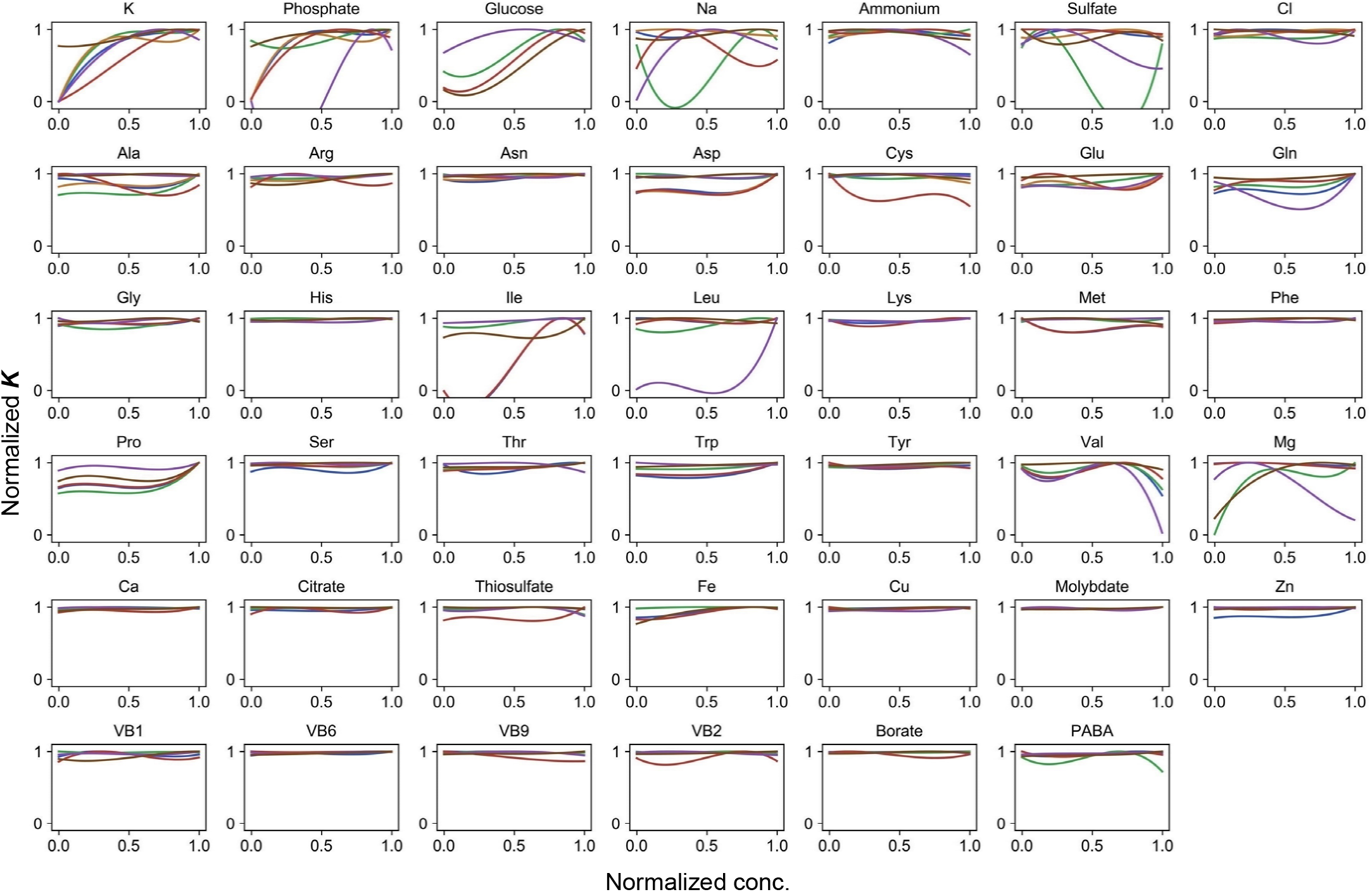
Normalized regression curves of the saturated population size. Colour variation indicates the alternative combinations of the other 40 components.

**Figure S6.**
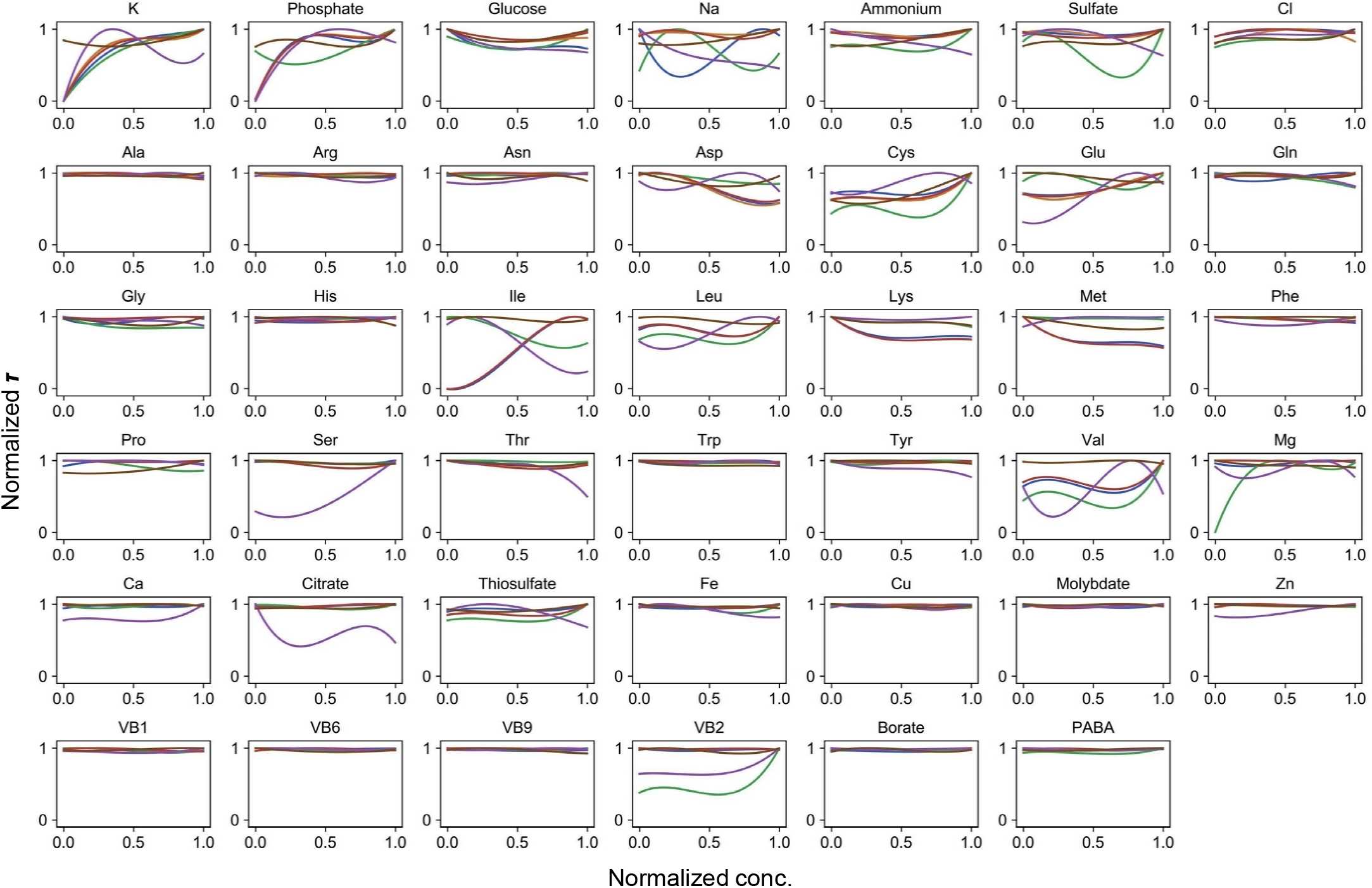
Normalized regression curves of the lag time. Colour variation indicates the alternative combinations of the other 40 components.

**Figure S7.**
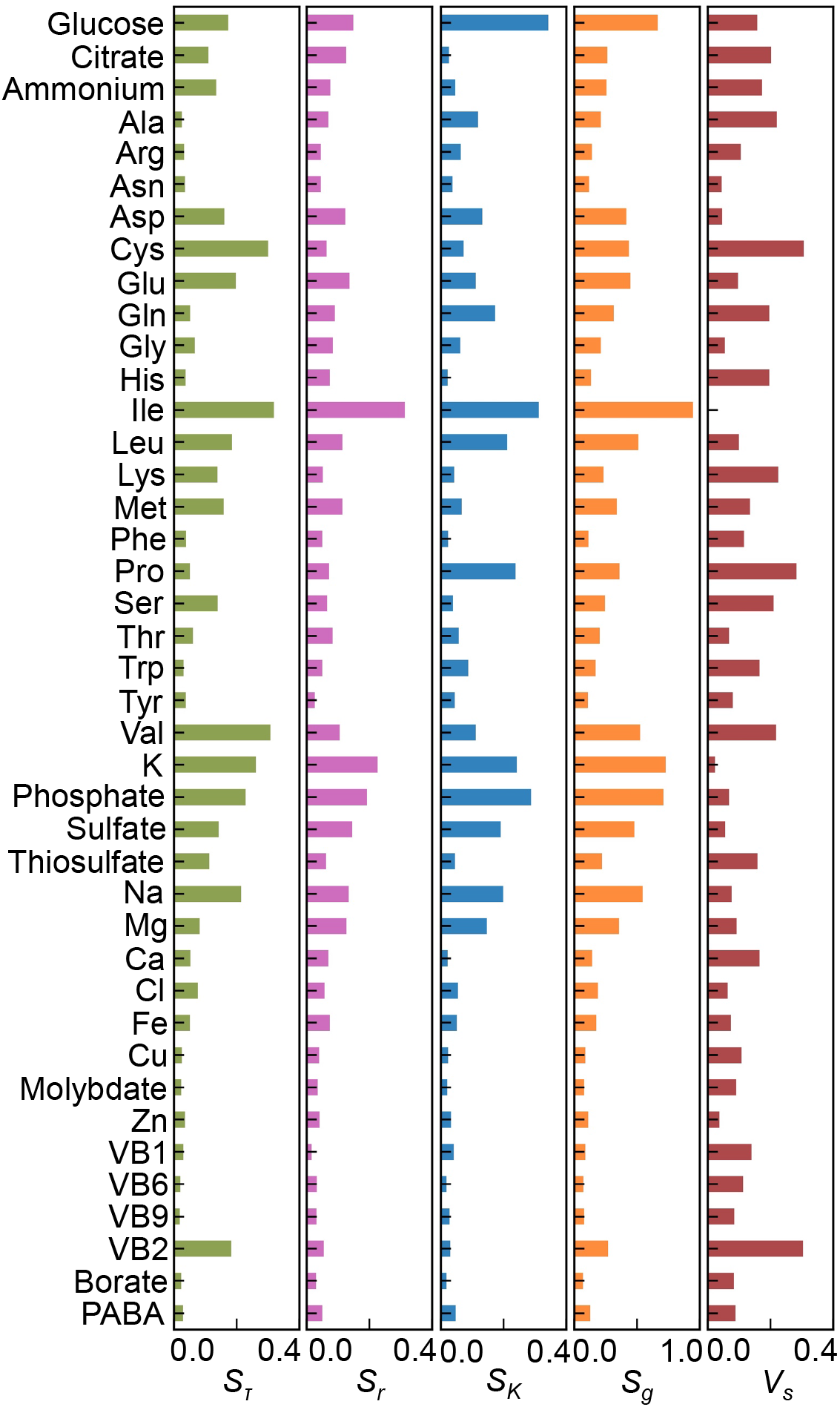
Sensitivity of the components. The values of *S*_*τ*_, *S*_*r*_, *S*_*K*_, *S*_*g*_ and *V*_*s*_ are shown.

**Figure S8.**
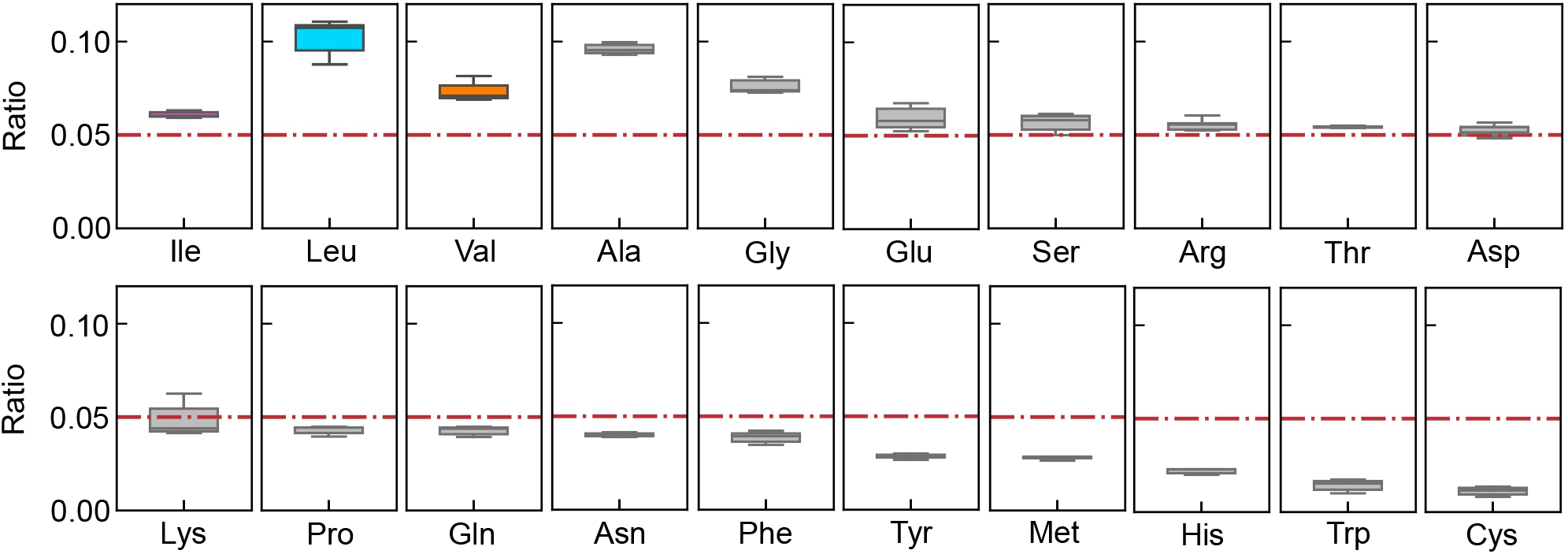
Relative abundance of 20 amino acids in growing *E. coli*. The boxplots represent the relative ratios estimated according to the genome and transcriptome information, and the red lines indicate the theoretical ratios of the 20 amino acids.

**Figure S9.**
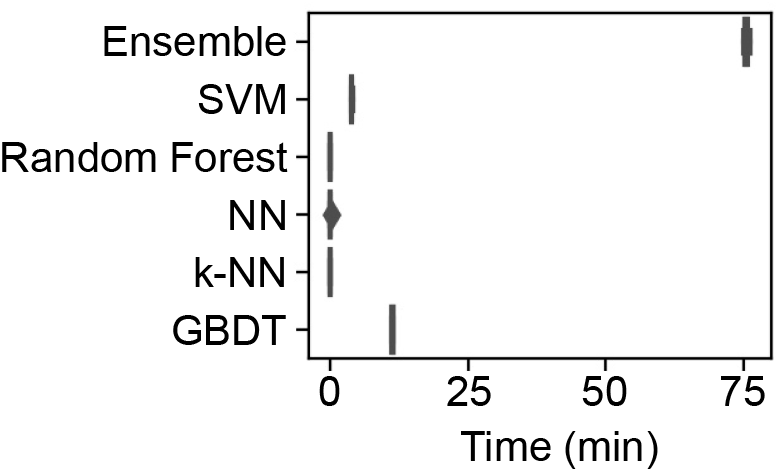
Time required for the ML model training. The time used to train the ML models by the supercomputer is shown in the boxplots of five independent replicates.

**Figure S10.**
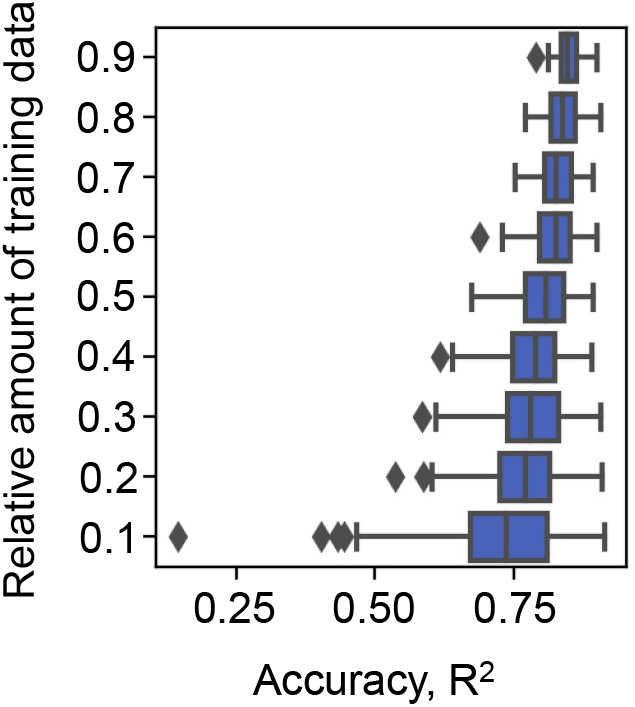
Effect of the abundance of training data on the accuracy of GBDT. Ten to ninety percent of the big data (the growth rate dataset) were randomly selected for model training. The accuracy of the growth rates predicted by the trained GBDT model was evaluated by R^2^. Model training by random selection and GBDT prediction were performed 100 times at each relative abundance of training data. The boxplots represent the accuracy of the 100 training and prediction runs of the growth rate by GBDT.

**Figure S11.**
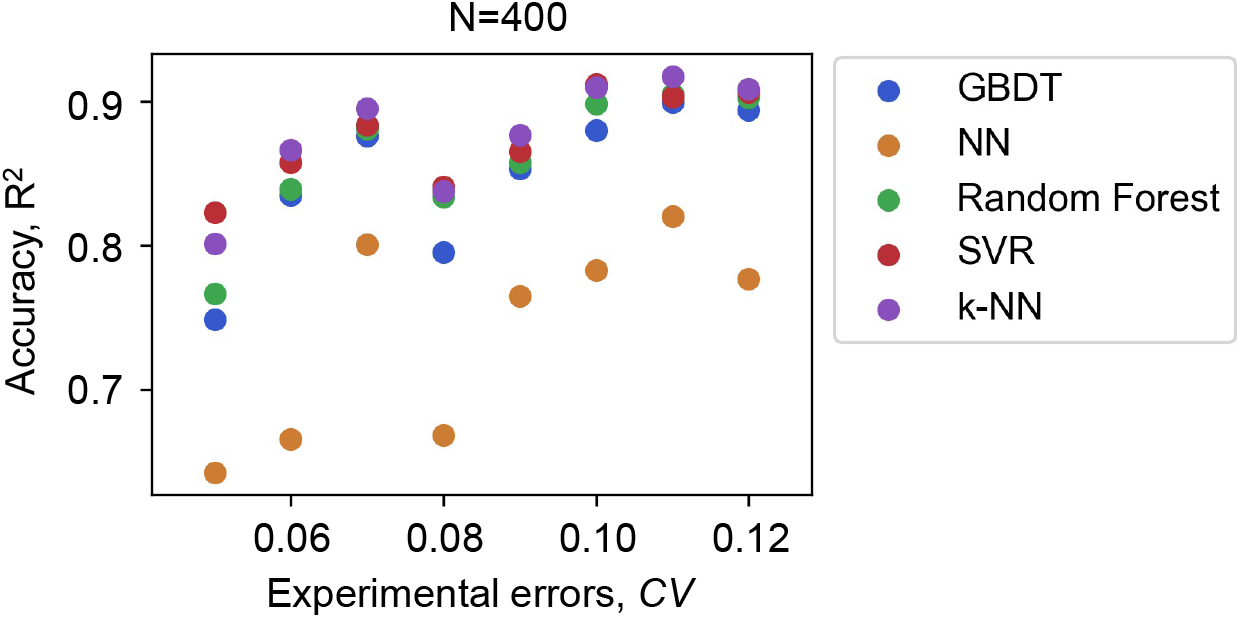
Accuracy of the ML models varied with the experimental errors of the data for training. Five ML models and the number of data points used in common are indicated. The experimental errors caused by the biological replications are shown as the coefficient of variance, *CV*.

**Figure S12.**
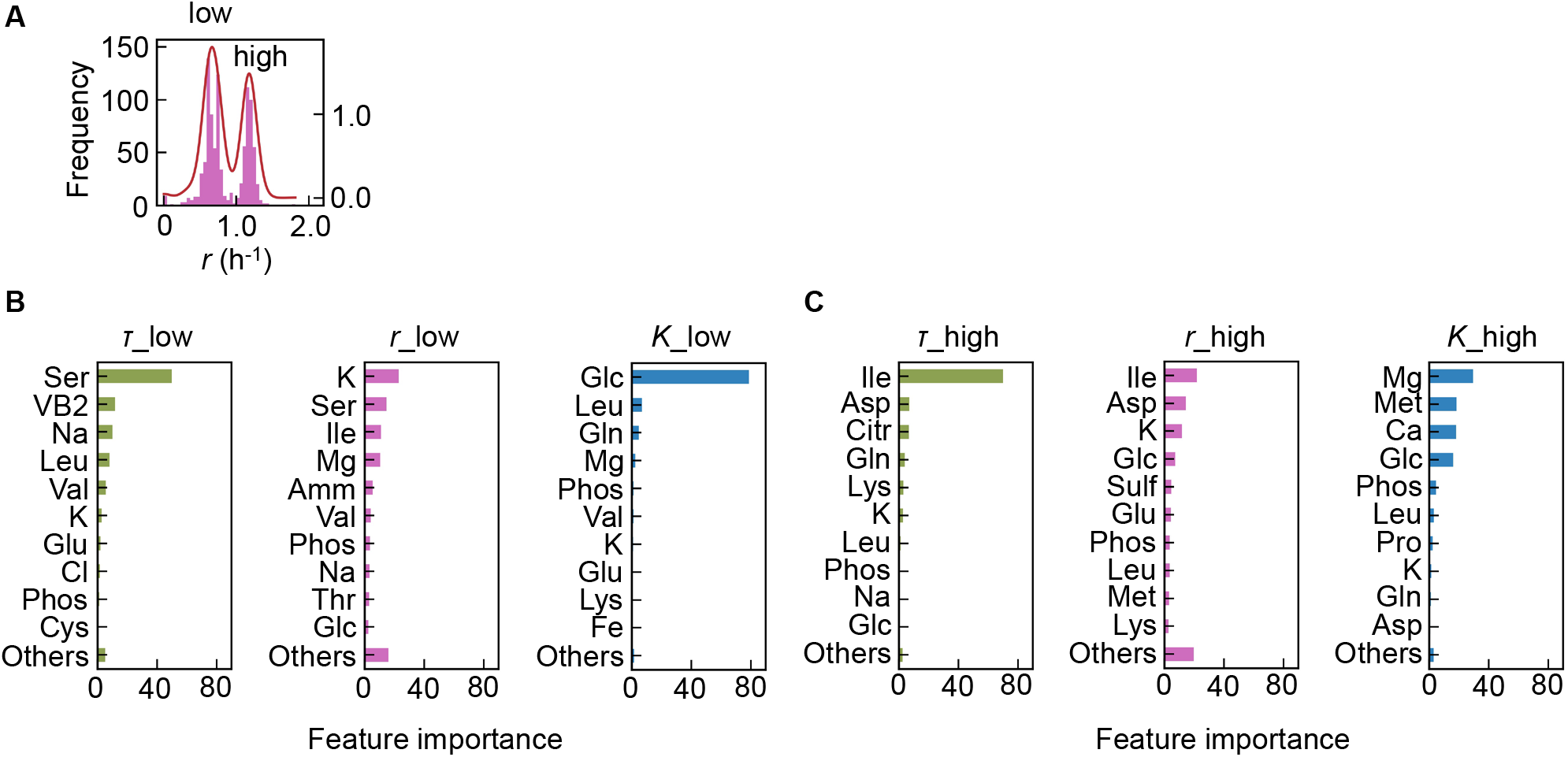
Separation of the multimodal distribution of *r*. The continuous probability distribution of the multimodal distribution of *r* (**A**) determined by Gaussian kernel density estimation is indicated by the red lines. The two separated distributions (datasets) are indicated as low and high. GBDT predictions of the low (**B**) and high (**C**) distributions are shown. Ten components with large contributions to the three parameters *τ*, *r*, and *K* are shown in order. The remaining 31 components are summed as “Others”.

**Figure S13.**
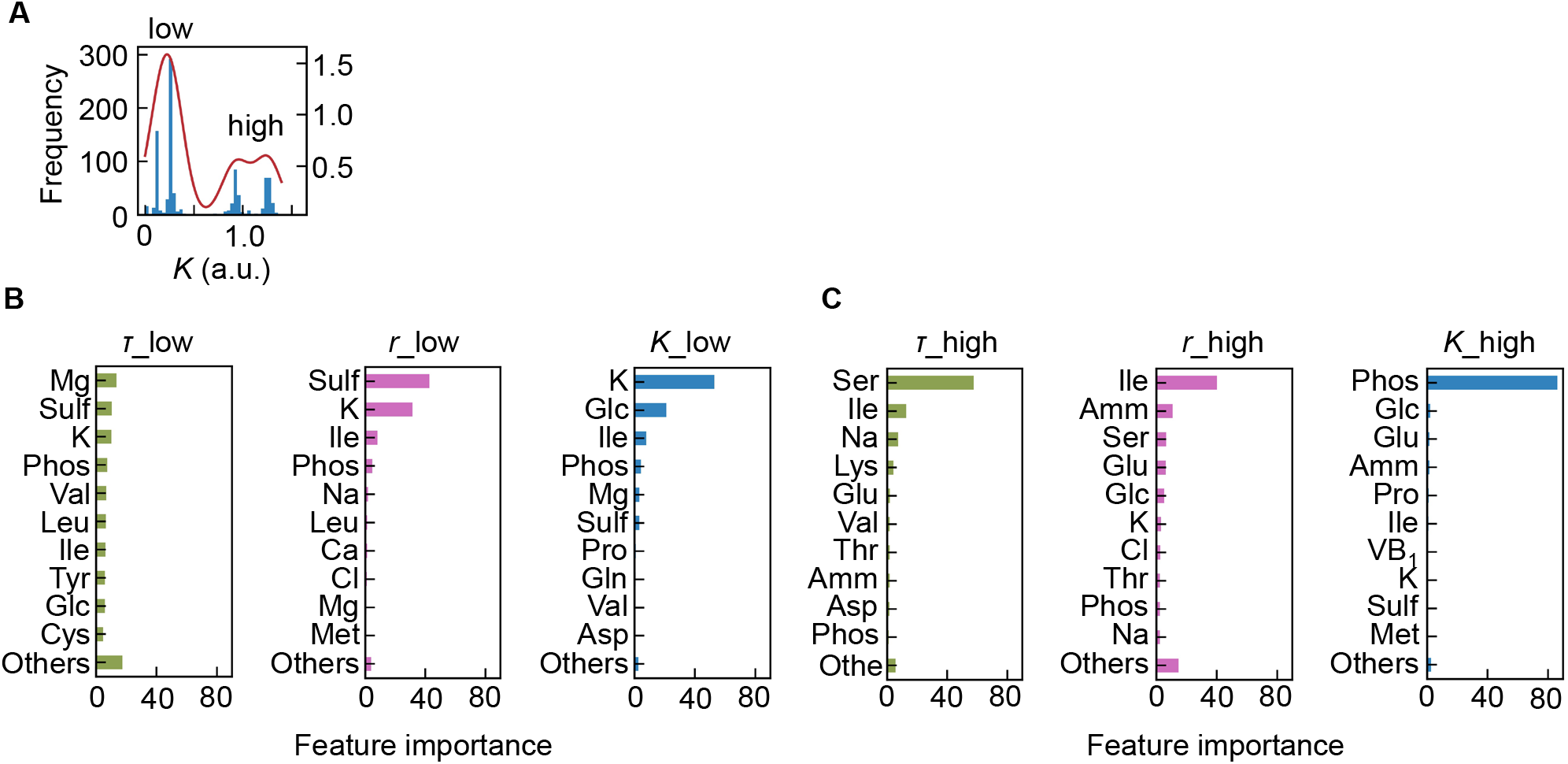
Separation of the multimodal distribution of *K*. The continuous probability distribution of the multimodal distribution of *K* (**A**) determined by Gaussian kernel density estimation is indicated by the red lines. The two separated distributions (datasets) are indicated as low and high. GBDT predictions of the low (**B**) and high (**C**) distributions are shown. Ten components with large contributions to the three parameters *τ*, *r*, and *K* are shown in order. The remaining 31 components are summed as “Others”.

**Figure S14.**
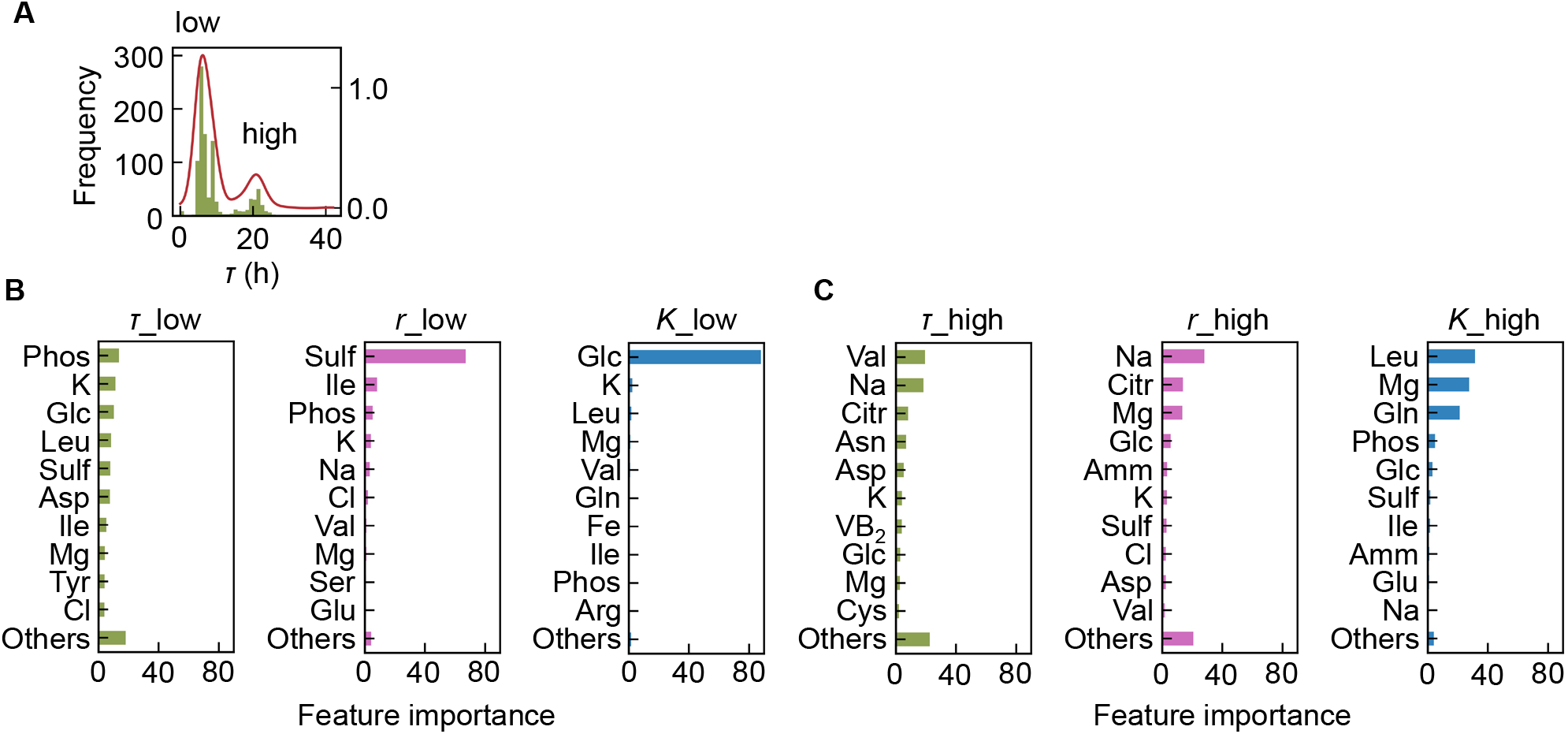
Separation of the multimodal distribution of *τ*. The continuous probability distribution of the multimodal distribution of *τ* (**A**) determined by Gaussian kernel density estimation is indicated by the red lines. The two separated distributions (datasets) are indicated as low and high. GBDT predictions of the low (**B**) and high (**C**) distributions are shown. Ten components with large contributions to the three parameters *τ*, *r*, and *K* are shown in order. The remaining 31 components are summed as “Others”.

**Figure S15.**
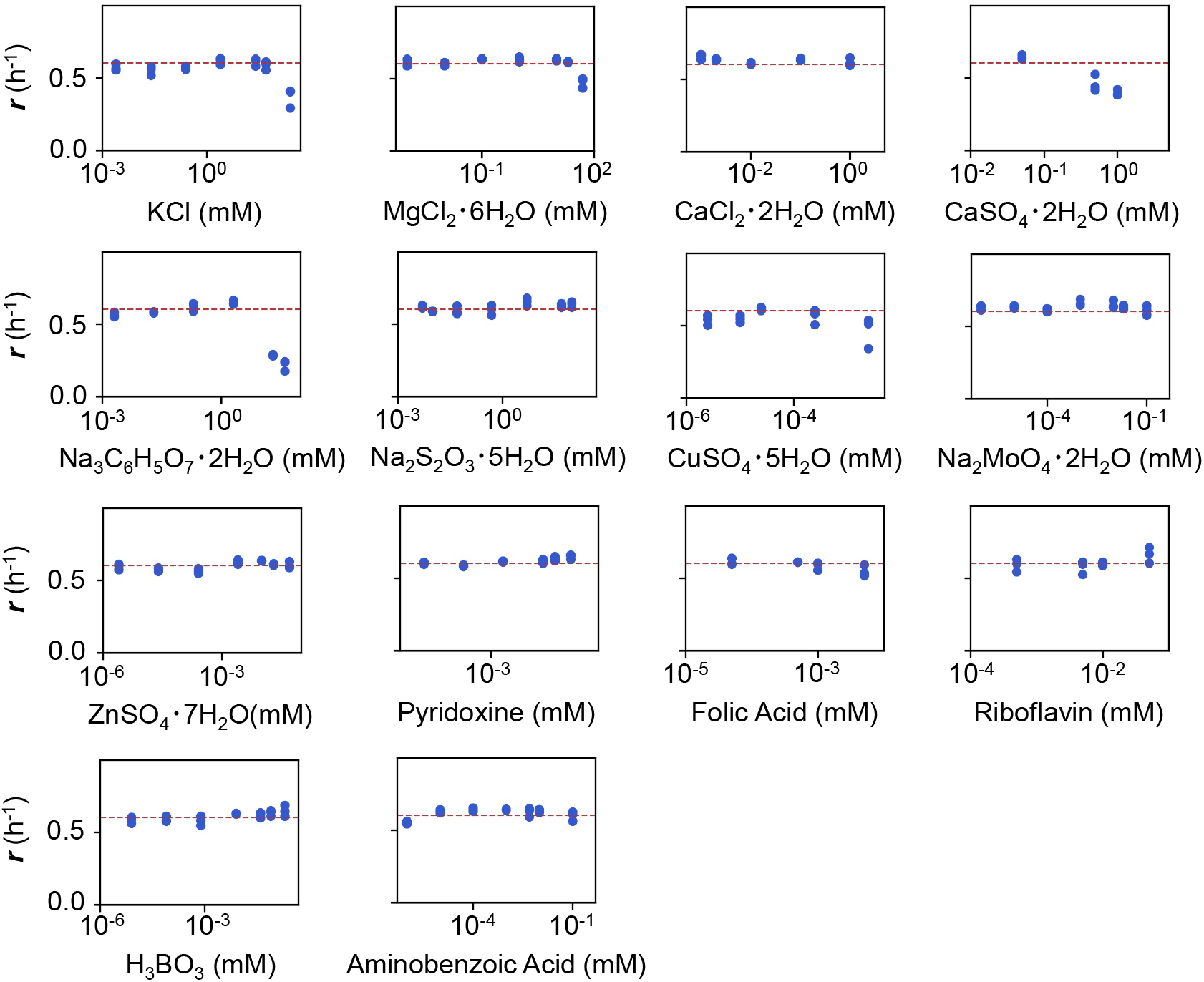
Experimental tests of the changes in *r* responding to the concentration gradients of the minor compounds. The product names of the compounds and the logarithmic concentrations are shown. The three biological replicates are shown as blue dots. The red lines indicate the condition of M63 minimal medium.

**Figure S16.**
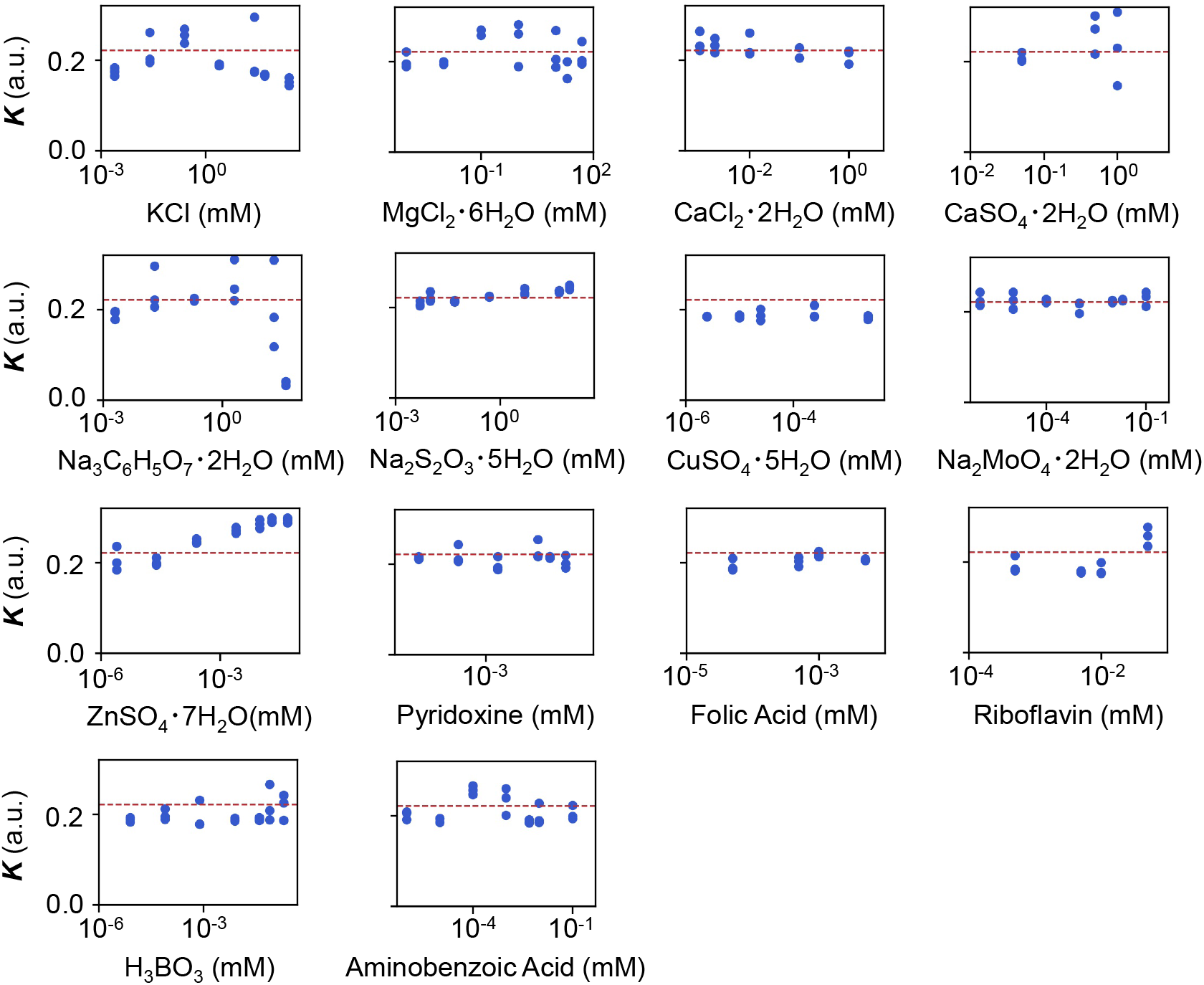
Experimental tests of the relationship between *K* and the minor compounds. The product names of the compounds and the logarithmic concentrations are shown. The three biological replicates are shown as blue dots. The red lines indicate the condition of M63 minimal medium.

